# Disruption of Mitochondrial Dynamics and Stasis Leads to Liver Injury and Tumorigenesis

**DOI:** 10.1101/2025.02.11.637688

**Authors:** Xiaowen Ma, Xiaoli Wei, Mengwei Niu, Chen Zhang, Zheyun Peng, Wanqing Liu, Junrong Yan, Xiaoyang Su, Shaolei Lu, Wei Cui, Hiromi Sesaki, Wei-Xing Zong, Hong-Min Ni, Wen-Xing Ding

## Abstract

**Background & Aims:** Mitochondrial dysfunction has been implicated in aging and various cancer development. As highly dynamic organelles, mitochondria constantly undergo fission, mediated by dynamin-related protein 1 (DRP1, gene name *Dnm1l*), and fusion, regulated by mitofusin 1 (MFN1), MFN2, and optic atrophy 1 (OPA1). However, whether and how dysregulation of mitochondria dynamics would be involved in liver pathogenesis and tumorigenesis is unknown.

**Methods:** *Dnm1l* Flox/Flox (*Dnm1l*^F/F^), *Mfn1*^F/F^ and *Mfn2*^F/F^ mice were crossed with albumin-Cre mice to generate liver-specific *Dnm1l* knockout (L-*Dnm1l* KO), L-*Mfn1* KO, L-*Mfn2* KO, L-*Mfn1, Mfn2* double KO (DKO), and L-*Mfn1, Mfn2, Dnm1l* triple KO (TKO) mice. These mice were housed for various periods up to 18 months. Some mice also received hydrodynamic tail vein injections of a Sleeping Beauty transposon-transposase plasmid system with *c-MYC* and *YAP*. Blood and liver tissues were harvested for biochemical and histological analysis.

**Results:** L-*Dnm1l* KO mice had elevated serum alanine aminotransferase levels and increased hepatic fibrosis as early as two months of age. By 12 to 18 months, male L-*Dnm1l* KO mice developed spontaneous liver tumors, primarily hepatocellular adenomas. While female L-*Dnm1l* KO mice also developed liver tumors, their incidence was much lower.

In contrast, neither L-*Mfn1* KO nor L-*Mfn2* KO mice had notable liver injury or tumorigenesis. However, a small portion of DKO mice developed tumors at 15-18 month-old. Increased DNA damage, senescence and compensatory proliferation were observed in L-*Dnm1l* KO mice but were less evident in L-*Mfn1* KO, L-*Mfn2* KO or DKO mice, indicating that mitochondrial fission is more important to maintain hepatocyte homeostasis and prevent liver tumorigenesis. Interestingly, further deletion of *Mfn1* and *Mfn2* in L-*Dnm1l* KO mice markedly abolished liver injury, fibrosis, and both spontaneous and oncogene-induced tumorigenesis. RNA sequencing and metabolomics analysis revealed significant activation of the cGAS-STING-interferon pathway and alterations in the tumor microenvironment pathways, alongside increased pyrimidine synthesis and metabolism in the livers of L-*Dnm1l* KO mice. Notably, the changes in gene expression and pyrimidine metabolism were considerably corrected in the TKO mice.

**Conclusions:** Mitochondrial dynamics and stability are essential for maintaining hepatic mitochondrial homeostasis and hepatocyte functions. Loss of hepatic DRP1 promotes liver tumorigenesis by increasing pyrimidine metabolism and activating the cGAS-STING-mediated innate immune response.

## Introduction

Primary liver cancer is the sixth most prevalent cancer and the third leading cause of global cancer mortality as of 2020, with an estimated 906,000 new cases and 830,000 deaths each year (Sung, Ferlay et al. 2021). Hepatocellular carcinoma (HCC) is the most common type of primary liver cancer that accounts for 75%-85% of instances and has an increasing incidence over the years (Sung, Ferlay et al. 2021). Although early HCC is curable by surgical resection, in most cases, HCC is diagnosed at an advanced stage, from which the standard anti-angiogenic drugs, such as sorafenib and lenvatinib, can only slightly improve the survival rate (Llovet, Ricci et al. 2008). Identification of novel therapeutic strategies is thus urgently needed for this challenging global health concern.

Recent studies have shown that mitochondria in HCC tumor tissues exhibit distinct morphological features, which are associated with HCC cell survival and promotes the infiltration of tumor-associated macrophages (Huang, Zhan et al. 2016, Bao, Zhao et al. 2019). Mitochondria size is strictly regulated by fission/fusion machinery: mitofusin 1 (MFN1), MFN2, and optic atrophy 1 (OPA1) mediate the fusion of mitochondrial outer and inner membranes, while dynamin-related protein 1 (DRP1) (gene name *DNM1L*) serves as major fission protein, driving mitochondrial fragmentation (Ni, Williams et al. 2015, Ma, Niu et al. 2024). Mitochondrial fusion is widely recognized for maintaining the fidelity of mitochondrial DNA (mtDNA), whereas dysfunctional mitochondria normally undergo fragmentation and tend to be removed by selective mitophagy and replaced via mitochondria biogenesis (Ni, Williams et al. 2015, Ma, Niu et al. 2024). In addition to quality control, mitochondrial morphological homeostasis is also closely involved in various stress-induced cellular responses, including apoptosis, energy transmission, innate and sterile inflammation, cell cycle regulation and dedifferentiation (Ma, Ni et al. 2024, Ma, Niu et al. 2024). Although clinical evidence regarding mitochondrial size in HCC tumor tissues is controversial, increased levels of fission protein DRP1 and decreased levels of fusion protein MFNs have been observed in both human HCC and mouse models of liver cancer (Wang, Lu et al. 2012, Lin, Qiu et al. 2020, Zhang, Li et al. 2020, Wang, Tian et al. 2022). Increased mitochondrial fission has been associated with a higher risk of distant metastasis and poor prognosis in HCC patients (Huang, Zhan et al. 2016, Lin, Qiu et al. 2020). However, it remains unclear whether the imbalanced fission and fusion events in cancer tissues represents an adaptive metabolic response for survival or actively contributes to HCC tumorigenesis.

Chronic liver injury is often associated with anomalous mitochondrial morphology (Bruguera, Bertran et al. 1977, Ma, McKeen et al. 2020, Yamada, Murata et al. 2022), suggesting dysregulated mitochondrial dynamics in precancerous conditions. From an etiological perspective, metabolic dysfunction-associated steatotic liver disease (MAFLD) and alcohol-associated liver disease (ALD) have emerged as major contributors to HCC development, and alcohol-associated cirrhosis is now the second-most common cause of HCC worldwide, following hepatitis B virus (HBV) infection (Akinyemiju, Abera et al. 2017). Both MAFLD and ALD have been shown to induce hepatic megamitochondria formation (Yamada, Murata et al. 2022, Ma, Chen et al. 2023). In ALD, megamitochondria formation is initially a respiratory adaptive response to facilitate alcohol metabolism, but accumulates under chronic alcohol exposure, leading to mitochondrial maladaptation and liver injury (Ma, Chen et al. 2023). Whether the long-standing megamitochondria also contribute to liver tumor formation is unknown. This study investigates the role of mitochondria fission/fusion homeostasis in liver tumorigenesis by generating genetically engineered mice with impaired mitochondrial fission (*Dnm1l*) and/or fusion (*Mfn1* or *Mfn2*).

## Material and Methods

### Animal experiments

*Dnm1l* Flox/Flox ^(F/F)^ mice (C57BL/6/129), *Mfn1,Mfn2* Flox/Flox ^(F/F)^ mice (C57BL/6/129) and *Dnm1l,Mfn1,Mfn2* Flox/Flox ^(F/F)^ mice (C57BL/6/129) were generated as described previously (Kageyama, Hoshijima et al. 2014) and were crossed with albumin-Cre mice (Alb-Cre, C57BL/6J) (Jackson Laboratory) for six generations. All animals were specific pathogen-free and maintained in a barrier rodent facility under standard experimental conditions. All procedures were approved by the Institutional Animal Care and Use Committee of the University of Kansas Medical Center. Blood and liver tissues were collected at indicated mouse age. Liver injury was determined by measuring serum alanine aminotransferase (ALT) and aspartate aminotransferase (AST). Liver sections and hematoxylin and eosin (H&E) staining were performed as described previously (Ni, Bockus et al. 2012).

### Hydrodynamic injection mouse model

Eight-week-old male L-*Dnm1l* KO, TKO and matched wild type (WT) mice were used in this study. Hydrodynamic injection was performed as previously described (Chen and Calvisi 2014). Briefly, 20 μg of pT3-EF1aH *c-MYC* and pT3-EF1aH *YAP S127A* along with 1.6 μg pCMV-SB10 transposase at a ratio of 25:1 was diluted in Ringer’s solution at a volume equivalent to 10% of the mouse’s body weight (g/mL). The solution was filtered through a 0.22 μm filter (Millipore, GSWP04700), and injected into the lateral tail vein in 5–7 seconds. Liver tissues and blood were collected 8 weeks post-injection.

### RNA extraction and RNA amplification

Total RNA was extracted from mouse livers using TRIzol reagent (15596-026; Ambion, ThermoFisher Scientific). For RNA sequencing analysis, total RNA were assessed for quality using a Qubit assay for quantification and an Agilent TapeStation gel analysis for integrity. Sequencing libraries were constructed using 1 µg of total RNA using the NEBNext Ultra II Directional RNA Library Prep Kit for Illumina (NEB). The sequencing library construction process includes mRNA purification by the polyA tails with polyT magnetic beads, fragmentation, strand-specific cDNA synthesis, end repair, 3’ end adenylation, adapter ligation, and PCR amplification. The constructed sequencing libraries were validated and quantified with Qubit and TapeStation assays. The sequencing library preps were pooled together equally by ng amount and the nM concentration of the pool was verified with an Illumina KAPA Library Quantification qPCR assay (Roche). Each library was indexed with a barcode sequence and sequenced in a multiplexed fashion. An Illumina NextSeq 550 system was used to generate single-end, 75-base sequence reads from the libraries. Base calling was carried out by the instrument’s Real-Time Analysis (RTA) software. The base call files (bcl files) were demultiplexed and converted to compressed FASTQ files by bcl2fastq2.

### RNAseq data analysis

RNAseq data were analyzed using our previously established technical pipeline (Rom, Xu et al. 2019, Rom, Liu et al. 2020). Briefly, we used HISAT2 v.2.1.0.13 to map the high-quality reads to the mouse reference genome (GRCm38.90). Quantification of the gene expression was generated with HTSeq-counts v0.6.0. Significant differentially expressed genes (DEGs) were generated with the R package DESeq2. Statistical significance was calculated by adjusting the P values with the Benjamini-Hochberg’s false discovery rate (FDR) (Hochberg and Benjamini 1990). The clustered heatmaps of the cGAS-STING pathway genes were plotted with ClustVis (Metsalu and Vilo 2015). Principle component analysis (PCA) and volcano plots were prepared with R. Pathway analysis were performed by using Ingenuity Pathway Analysis (IPA). We analyzed the enrichment of up-and down-regulated genes among DRP1 KO and TKO group as compared to the WT group. Genes with a log2 fold change >1.5 or <-1.5 with an FDR-corrected q value <0.05 were used to perform the analysis.

### Liver Metabolomics Analysis

Metabolomics analysis was performed by Metabolon, resulting in a dataset comprising a total of 907 compounds of known identity (biochemicals). After log transformation and imputation of missing values using the minimum observed value for each compound, Welch’s two-sample t-test was applied to identify biochemicals that significantly differed between groups. A summary of the numbers of biochemicals that achieved statistical significance (p≤ 0.05) and those approaching significance (0.05 < p < 0.10) is provided.

### Liquid chromatography–mass spectrometry (LC-MS)

Twenty to forty milligrams of frozen liver tissue samples were weighed and kept in prechilled 2 mL Eppendorf tubes with beads and ground with CryoMill (Retsch NC1216630) at liquid nitrogen temperature and 20 Hz for 2 min. The ground samples were extracted by adding 40 times -20°C 40:40:20 methanol:acetonitrile:water solution with 0.1M formic acid, followed by vortexing and centrifugation at 16,000 x g for 10 min at 4°C. Forty-four microliters of 15% (m/v) NH4HCO3 was added to 500 μl of the transferred extract to neutralize the acid. The samples were centrifuged at 16,000 x g for 10 min to remove the protein precipitate. Then, 300 μl of the supernatant was collected into a new tube for LC-MS. LC-MS was performed at the Rutgers Cancer Institute Metabolomics Shared Resource, as previously described (Dai, Shen et al. 2022). The metabolite features were extracted in El-MAVEN with a mass accuracy window of 5 ppm. The metabolite annotation is based on accurate mass and retention time that matches the in-house library.

### Western blot analysis

Total liver lysates were prepared using radioimmunoprecipitation assay (RIPA) buffer (1% NP40, 0.5% sodium deoxycholate, 0.1% sodium dodecyl (lauryl) sulfate). Proteins (30μg) were separated by 8%–12% SDS-PAGE gel before transfer to a PVDF membrane. Membranes were probed using indicated primary and secondary antibodies and developed with SuperSignal West Pico Plus Chemiluminescent Substrate (Thermo scientific, 34579) and Immobilon Western Chemiluminescent HRP Substrate (Millipore, WBKLS0500). All the primary and secondary antibodies used in this study were listed in **Table 1**.

**Table 1.** List of antibodies used for western blot, and IHC staining.

| Antibodies | Brand | Catalogue Number |
| --- | --- | --- |
| 4EBP1 | Cell Signaling | 9452 |
| p-4EBP1 (Ser65) | Cell Signaling | 9451 |
| AKT (pan) | Cell Signaling | 4691 |
| p-AKT (Ser473) | Cell Signaling | 4058 |
| α-SMA | Abcam | Ab5694 |
| β-actin | Sigma | A5441 |
| BAX | Santa Cruz | sc-493 |
| β-Catenin | Cell Signaling | 9581 |
| BIP | Sigma | G9043 |
| CD133 | Proteintech | 18470-1-AP |
| cGAS | Cell Signaling | 31659 |
| CHOP | Santa Cruz | Sc-793 |
| Cleaved Caspase3 | Cell Signaling | 9661 |
| cMYC | Gene Tex | GTX103436 |
| DRP1BD Bioscience611112ERK | Cell Signaling | 9102 |
| p-ERK (Thr202/Tyr204) | Cell Signaling | 9101 |
| F4/80 | Invitrogen | 14-4801 |
| GAPDH | Cell Signaling | 2118 |
| GCLM | Gift from Dr. Terry Kavanagh's lab at University of Washington, Seattle, WA |  |
| Glypican3 | Abcam | Ab66596 |
| GSK-3α/β | Cell Signaling | 5676 |
| p-GSK3β (Ser9) | Cell Signaling | 5558 |
| IRF3 | Cell Signaling | 4302 |
| IRF7 | Cell Signaling | 72073 |
| Ki67 | Gene Tex | GTX16667 |
| Mfn1 | Santa Cruz | sc-50330 |
| Mfn2 | Sigma | M6444 |
| Myeloperoxidase (MPO) | BioCare medical | PP023AA |
| MST1 | Cell Signaling | 3682 |
| NQO1 | Abcam | ab28947 |
| p21 | Santa Cruz | sc-6246 |
| p27 | Cell Signaling | 3698 |
| p62 | Abnova | H00008878-M01 |
| p-γ-H2AX (Ser139) | Cell Signaling | 9718 |
| PCNA | Santa Cruz | sc-56 |
| PTEN | Cell Signaling | 9552 |
| p-PTEN (Ser380/Thr382/383) | Cell Signaling | 9554 |
| S6 | Cell Signaling | 2217 |
| p-S6 (Ser235/236) | Cell Signaling | 4858 |
| SOX9 | Millipore | AB5535 |
| STING | Cell Signaling | 50494S |
| TAZ | Cell Signaling | 83669 |
| TBK1 | Cell Signaling | 3504 |
| p-TBK1(Ser172) | Cell Signaling | 5483 |
| YAP | Cell Signaling | 14074 |
| p-YAP (Ser127) | Cell Signaling | 4911 |
| HRP-Goat Anti-Mouse IgG | Jackson | 115-035-146 |
| HRP-Goat Anti-Rabbit IgG | Jackson | 111-035-144 |

### Electron microscopy

Fine-cut liver tissues were fixed with 2% glutaraldehyde in phosphate buffer 0.1 mol/L (pH 7.4) followed by 1% OsO4. After dehydration, thin sections were stained with uranyl acetate and lead citrate for observation under a JEM 1016CX electron microscope (JEOL, Tokyo, Japan). Images were acquired digitally.

### Histological staining

A standard immunohistochemistry procedure was applied in this study. Briefly, paraffin-embedded tissue sections were incubated with primary antibodies at 4°C overnight after deparaffinization and heat-induced antigen retrieval in citrate buffer. Sections were then washed and incubated with secondary antibodies for 1h at 37°C and developed using ImmPACT NovaRED HRP substrate (Vector Labs, SK-4805). Tissues were counterstained with hematoxylin. The number of IHC-positively stained cells was counted from 15 different fields (200X) in a double-blinded fashion. For Sirius Red staining, paraffin-embedded liver tissue sections were dewaxed and rehydrated followed by staining sections with Sirius Red Solution.

### β-Galactosidase (β-Gal) staining

Liver cryosections were used for β-Gal staining follow the manufacturer’s instruction (Cell Signaling Technology #9860). Briefly, liver cryosections were fixed with 1x Fixative Solution for 15 min at room temperature and rinsed with 1x PBS for 2 times. β-Galactosidase Staining Solution was applied for a 37°C overnight incubation until a desired color was reached. Tissue sections were imaged immediately thereafter.

### Statistical analysis

All experimental data were expressed as mean ± SE and subjected to one-way ANOVA analysis with Bonferroni post hoc test or Student’s t-test where appropriate. Mean ± SD were used for the image quantitative data. A p-value less than 0.05 was considered significant.

## Results

### L-*Dnm1l* KO mice have an increased number of megamitochondria, liver injury, fibrosis and spontaneous liver tumorigenesis, which can be rescued by restoration of mitochondrial stasis

L-*Dnm1l* KO, L-*Mfn1, Mfn2* DKO (DKO) and L-*Mfn1, Mfn2, Dnm1l* TKO (TKO) mice were born at the expected Mendelian ratio, and the appearance of newborns was normal. An approximately 5-fold increase in serum ALT and a 2-fold increase in serum AST activities were found in L-*Dnm1l* KO mice at 2-month-old (2M) compared to their matched WT littermates, which were completely rescued by further deletion of mitochondrial fusion proteins MFN1 and MFN2 **(Fig. 1A-B)**. L-*Dnm1l* KO mice had increased hepatic Sirus red staining and hepatic α-smooth muscle actin (α-SMA) levels than WT mice, which is blunted in TKO mice (**Fig. 1C & Supplemental Fig 1**). Ultrastructure electron microscopy (EM) analysis showed heterogeneity of mitochondrial size in hepatocytes of mouse liver, but the number of large-sized or elongated mitochondria was more than 2-fold higher in L-*Dnm1l* KO mice than in WT or TKO mice **(Fig. 1D-E)**. These data suggest that disruption of hepatic mitochondrial fission leads to increased hepatic megamitochondria and liver injury in mice, which can be rescued by restoring mitochondrial morphological homeostasis.

**Figure 1.**
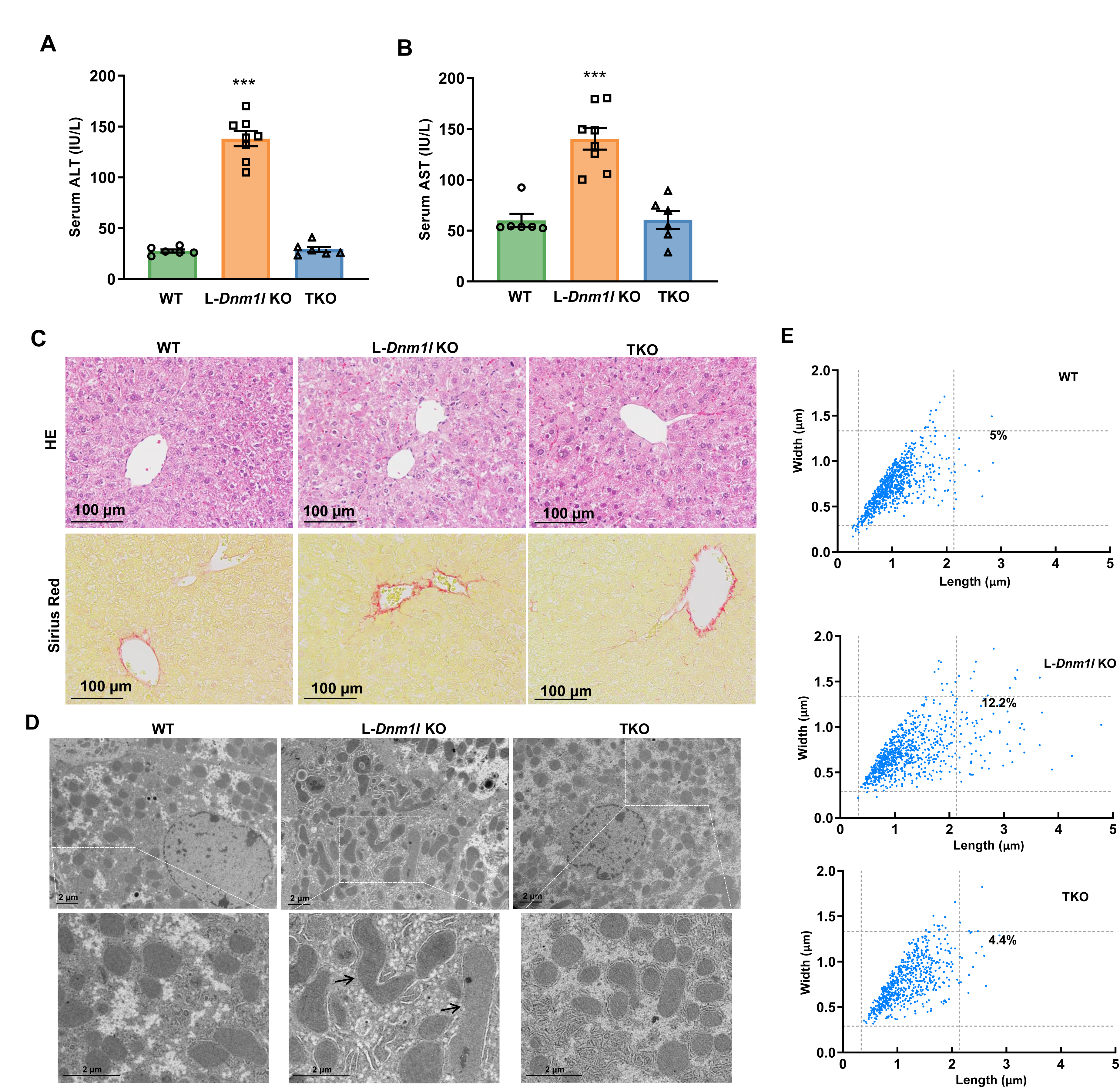
L-*Dnm1l* KO mice have an increased number of megamitochondria and liver injury, restored by additional *Mfn1, Mfn2* ablation. (A-B) Serum ALT and AST activities of 2M-old mice with indicated genotypes. (C) Representative H&E and Sirius Red staining from 2M-old mouse liver. (D) Representative EM images of the livers from 2M-old mice. (E) Quantification of mitochondrial size from EM images. The reference lines represent 95th percentile in WT group. 2-3 mice were quantified in each group (mitochondria number=600-900). All results are expressed as means± SEM (n=6-8). *p<0.05, **p<0.01, ***p<0.001; Student’s t-test compared to WT group.

Starting at the age of 12M, L-*Dnm1l* KO mice developed spontaneous liver tumors, with 64% incidence at 12M and 92% incidence after 15-18M in male mice. The incidence of liver tumors was much lower in female L-*Dnm1l* KO mice than in the age-matched L-*Dnm1l* KO male mice. Neither L-*Mfn1* KO nor L-*Mfn2* KO mice had an obvious liver injury or visible tumor formation **(Fig. 2A-D)**. However, a small portion (29%) of DKO mice developed liver tumors from 15M, but the tumor burden and maximum size were greatly lower than age-matched L-*Dnm1l* KO mice **(Fig. 2B-C)**. No visible tumors were found in the TKO mouse livers (**Fig. 2A-B)**, indicating that re-establishment of mitochondria stasis is critical to rescue the liver tumorigenesis induced by the unbalanced mitochondrial fission and fusion. The elevated serum ALT activity in L-*Dnm1l* KO mice persisted until 12M, with the peak in 2M, and normalized in 15-18M **(Fig. 1A, 2D)**. No significant changes in ALT activities were found in TKO or DKO mice **(Fig. 2D)**. Starting at 6M, the nuclei in L-*Dnm1l* KO hepatocytes showed a pleomorphic pattern with an increased number of vacuolated nuclei (**Fig. 2E, arrows**), which were more obvious in 12M L-*Dnm1l* KO hepatocytes but were absent in age-matched WT and TKO hepatocytes **(Fig. 2E)**. Moreover, we found severe liver fibrosis in L-*Dnm1l* KO mice, but not TKO or DKO, as early as 6M and persistent to 12M, showing prototypical perisinusoidal (chicken-wire) collagen fiber deposition by Sirius Red staining and increased hepatic α-SMA **(Fig. 2E & Supplemental Fig 1B-C)**, indicating increased wound-healing response to the chronic liver injury in L-*Dnm1l* KO mice. Taken together, impaired mitochondrial fission in hepatocytes leads to increased liver injury, fibrosis, and liver tumorigenesis in mice, which can be rescued with hepatic mitochondria stasis.

**Figure 2.**
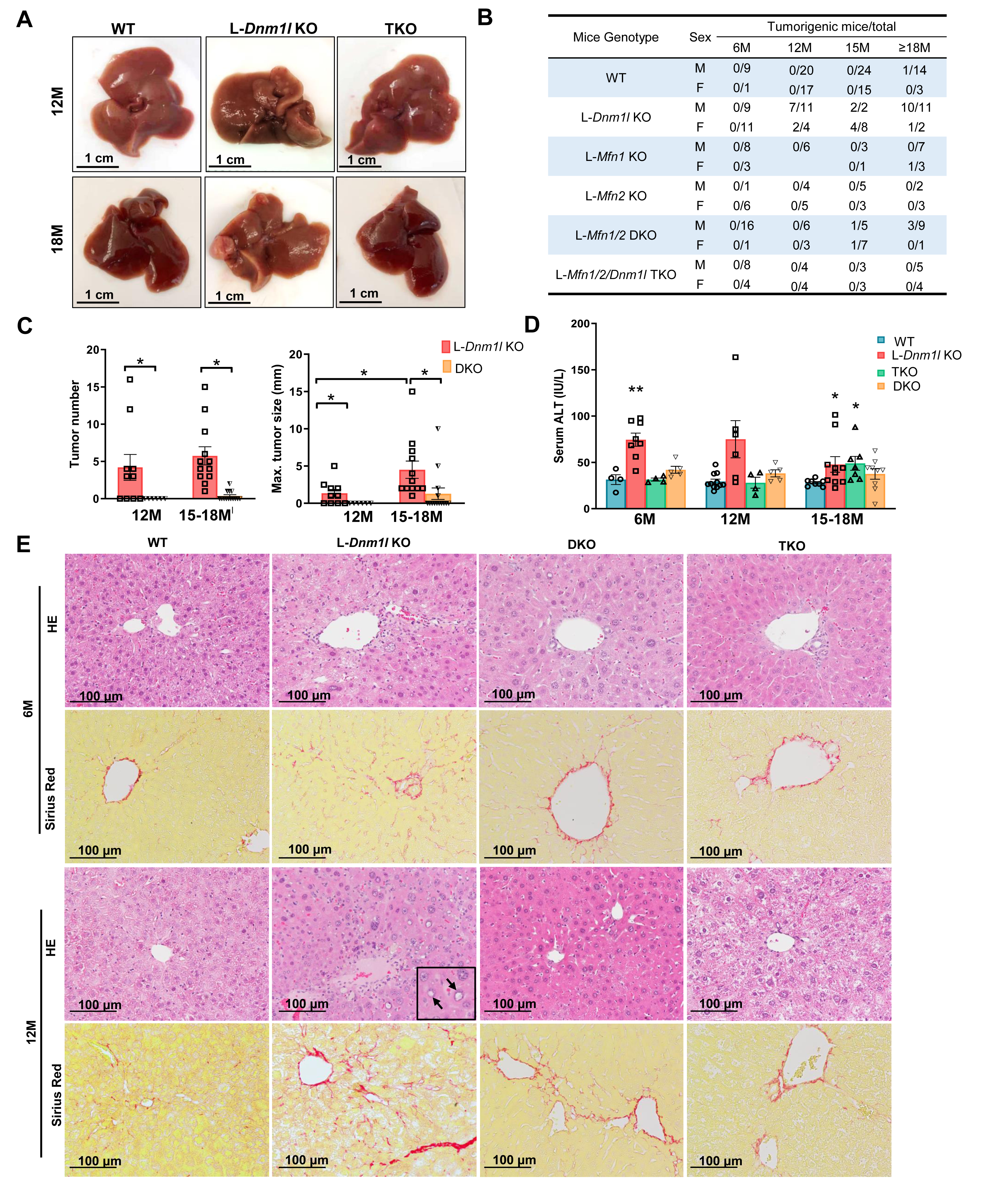
Spontaneous liver tumorigenesis and fibrosis in L-*Dnm1l* KO mice were rescued by further depletion of *Mfn1, Mfn2*. (A) Representative gross liver image. (B-C) Liver tumorigenesis in indicated genotyped mice. (D) Serum ALT activities. (E) Representative H&E and Sirius Red staining in the liver. Black arrows: vacuolated nuclei. All results are expressed as means± SEM (n=4-11 for ALT). *p<0.05, **p<0.01, ***p<0.001; One-Way ANOVA analysis with Bonferroni’s post hoc test for (C), Student’s t-test compared to corresponding WT group for (D).

### Loss of liver *Dnm1l* activates cGAS-STING-interferon pathway and induces liver inflammation, which are attenuated in TKO mouse livers

To investigate the potential mechanisms underlying early-stage liver injury in L-*Dnm1l* KO mice, we performed RNA sequencing analysis on liver samples from 2M and 6M mice. In 2M mice, principal component analysis (PCA) revealed a distinct separation in gene expression between the WT and L-*Dnm1l* KO groups, while the TKO group exhibited a gene expression profile closer to that of WT mice **(Fig. 3A)**. A similar gene distribution pattern was identified in 6M mice (**Supplemental Fig 2A**). In 2M mice, 743 up-regulated and 125 down-regulated genes were identified in L-*Dnm1l* KO mice compared to the WT, whereas only 48 differential genes were detected between the TKO and WT mice **(Fig. 3B)**. These findings suggest that the gene expression alterations in L-*Dnm1l* KO mice were effectively corrected by the reconstitution of “mitochondria stasis” through a simultaneous block of fusion and fission events in TKO mice. Compared to WT mice, many of the top up-regulated genes in L-*Dnm1l* KO mice, such as *Ifi44*, *Ifi27*, *Oasl2* and *Irf7*, belong to the cyclic GMP-AMP Synthase (cGAS)-stimulator of Interferon Genes (STING) - interferon innate immune pathway. In contrast, *Mfn1* and *Mfn2* are among the most significantly down-regulated genes in TKO mouse livers (**Fig. 3B**), confirming a successful deletion of these genes in TKO mice. Ingenuity Pathway Analysis revealed a strong activation of the ‘inflammation-related’ and ‘tumor microenvironment’ pathways in L-*Dnm1l* KO mice **(Fig. 3C)**, along with a notable upregulation of genes within the cGAS-STING-interferon pathway, as shown in the heatmap **(Fig. 3D)**. Interestingly, the increased expression of cGAS-STING-interferon pathway genes in L-*Dnm1l* KO mice persisted from 2M to 6M but was nearly completely abolished in TKO mice (**Fig. 3D, Supplemental Fig. 2B**). In addition, no significant changes in the expression of cGAS-STING-interferon pathway genes were observed in DKO mice compared to WT mice (**Supplemental Fig. 2B**), suggesting that inhibiting mitochondria fission, rather than fusion, is more critical for the activation of cGAS-STING innate immune pathway.

**Figure 3.**
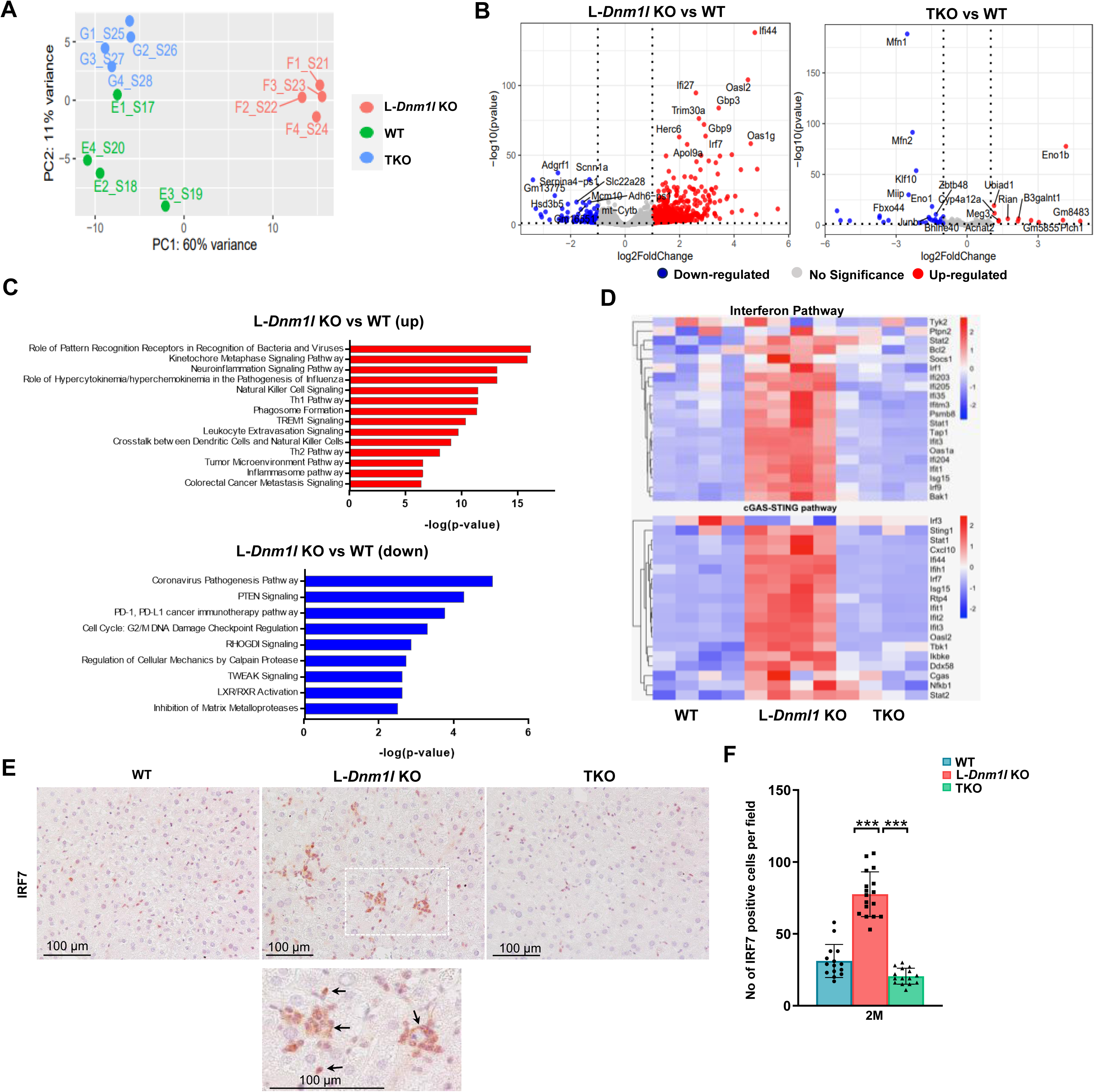
Loss of liver *Dnm1l* activates cGAS-STING-interferon pathway and promotes tumor microenvironment transformation, blunted in TKO mice. (A) Principal component analysis (PCA) of RNA-seq dataset. (B) Volcano plot of RNA-seq dataset. (C) The top up- and down-regulated pathways identified by Ingenuity Pathway Analysis (IPA) from RNA-seq dataset of indicated mouse liver tissues. (D) Heatmap of interferon and cGAS-STING pathway involved genes from the RNA-seq dataset. (E-F) Representative immunohistochemistry staining of IRF7 in 2M-old mouse liver. At least 5 fields were quantified from each mouse (n=3 mice). All results are expressed as means± SD. *p<0.05, **p<0.01, ***p<0.001; One-Way ANOVA analysis with Bonferroni’s post hoc test.

In parallel, IHC staining revealed enriched IRF7-positive immune cell infiltration in the livers of 2M L-*Dnm1l* KO mice **(Fig. 3E-F)**. Protein levels of cGAS, STING, p-TBK1, IRF3, and IRF7 were significantly elevated in the normal tissues of 15-18M L-*Dnm1l* KO mice, with even higher levels detected in the tumor tissues. However, most of these increases were reduced in age-matched TKO mice **(Supplemental Fig. 2C)**. Western blot analysis confirmed the successful deletion of hepatic DRP1, MFN1 and MFN2 in the KO, TKO and DKO mice (**Supplemental Fig. 2D**). In L-*Dnm1l* KO mouse liver, we found an increased numbers of F4/80+ macrophages and MPO+ neutrophils compared to WT and TKO mice. This phenomenon was particularly obvious in 2M mice but was attenuated by 6M **(Fig. 4A-B)**. Consistent with the IHC staining, RNAseq analysis indicated decreased expression of gene signatures in hepatocytes and increased expression in hepatic stellate cells (HSC) and Kupffer cells/macrophages in 2M and 6M L-*Dnm1l* KO mice, although the changes were less prominent in 6M. Interestingly, regardless of age, the alterations in cell-type-specific gene signatures were significantly blunted in TKO mice (**Supplemental Fig. 3A-B**). Furthermore, the levels of cleaved caspase-3 and endoplasmic reticulum (ER) stress markers, including BIP and CHOP, were markedly increased in the livers of 2M L- *Dnm1l* KO mice **(Fig. 4C)**, suggesting that hepatic DRP1 loss may induce ER stress, apoptosis and inflammation, which may contribute to the early-stage of liver injury in mice.

**Figure 4.**
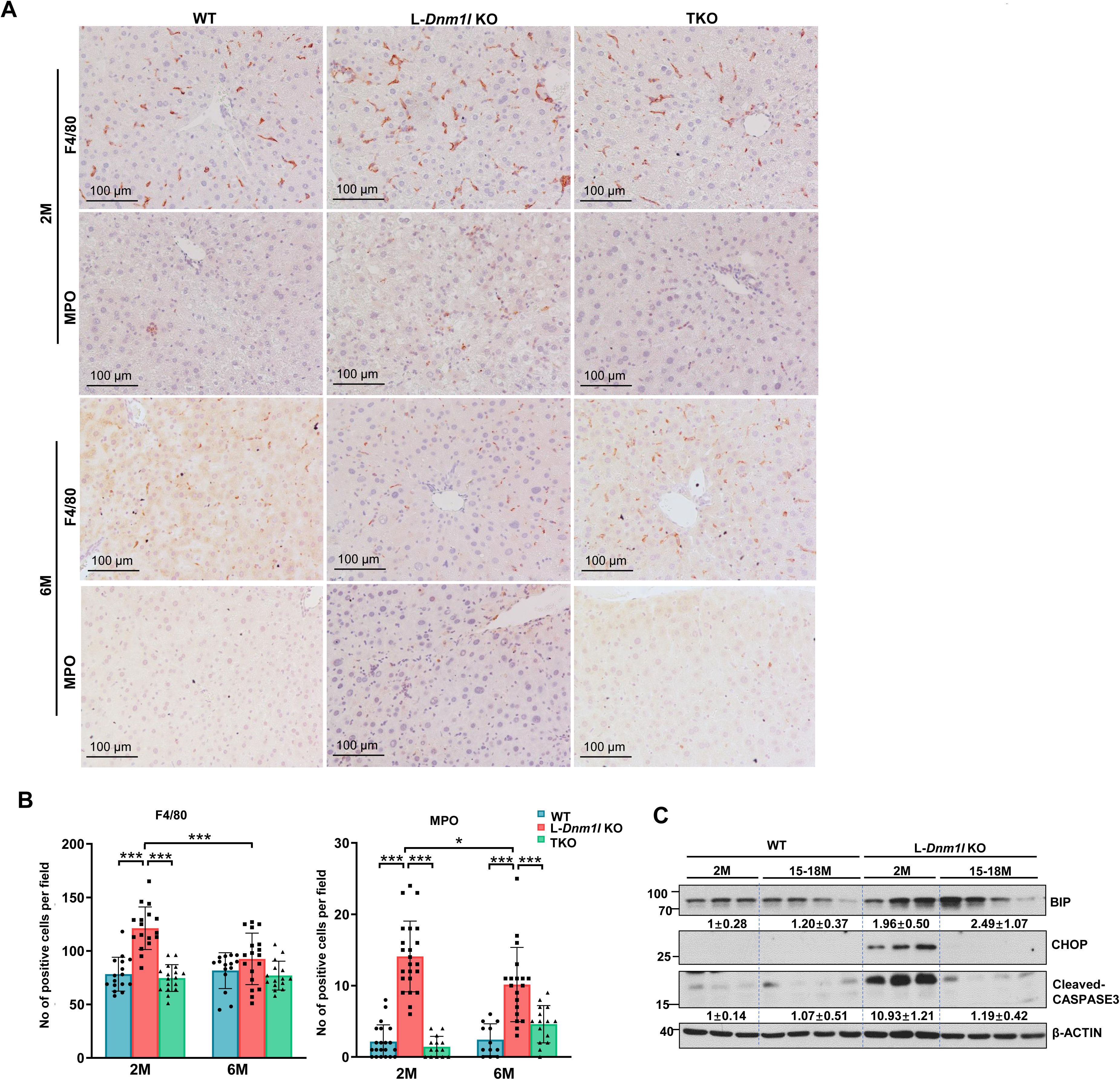
Loss of liver *Dnm1l* promotes endoplasmic reticulum (ER) stress and immune cell infiltration in mouse livers, which are suppressed in TKO mice. (A-B) Representative immunohistochemistry staining of F4/80 and MPO from indicated mouse liver. At least 5 fields were quantified from each mouse (n=3 mice). (C) Western blot analysis of total liver lysate. All results are expressed as means± SD. *p<0.05, **p<0.01, ***p<0.001; One-Way ANOVA analysis with Bonferroni’s post hoc test.

### Enhanced pyrimidine synthesis and decreased mitochondrial gene expression in L-*Dnm1l* KO mouse livers are restored in TKO mice

Metabolomics analysis of the mitochondrial TCA cycle, amino acids and nucleotide showed that pyrimidine-related metabolites, including glutamate, orotate, N-carbamoyl-aspartate, and uracil, exhibited the most significant increases in the livers of 2M L-Dnm1l KO mice compared to WT, DKO, and TKO mice (**Fig. 5A**). This trend was similarly observed in the livers of 6M mice (**Fig. 5B**). To rigorously validate the changes in pyrimidine metabolism, we conducted untargeted metabolomics analysis using a different batch of 2M mouse liver tissues, and the results confirmed an increase in pyrimidine metabolites, specifically dihydroorotate and orotate, which were elevated approximately 50-fold and 14-fold respectively, in the livers of L-*Dnm1l* KO mice compared to WT mice (**Fig 5C-D**). Dihydroorotate dehydrogenase (DHODH) is the mitochondrial enzyme in the *de novo* pyrimidine synthesis pathway that catalyzes the ubiquinone-dependent oxidation of dihydroorotate to orotate. Therefore, these results provide direct evidence that mitochondrial morphology affects pyrimidine metabolism. In contrast, no significant differences were observed in DKO or TKO mice relative to WT mice (**Figure 5C-D**). Consistent with the metabolomics data, RNA sequencing analysis revealed increased expression of genes related to pyrimidine metabolism in 2M L-*Dnm1l* KO mice compared to WT mice. These changes were less pronounced in 6M mice, as well as in TKO and DKO mice (**Fig 5E**).

**Figure 5.**
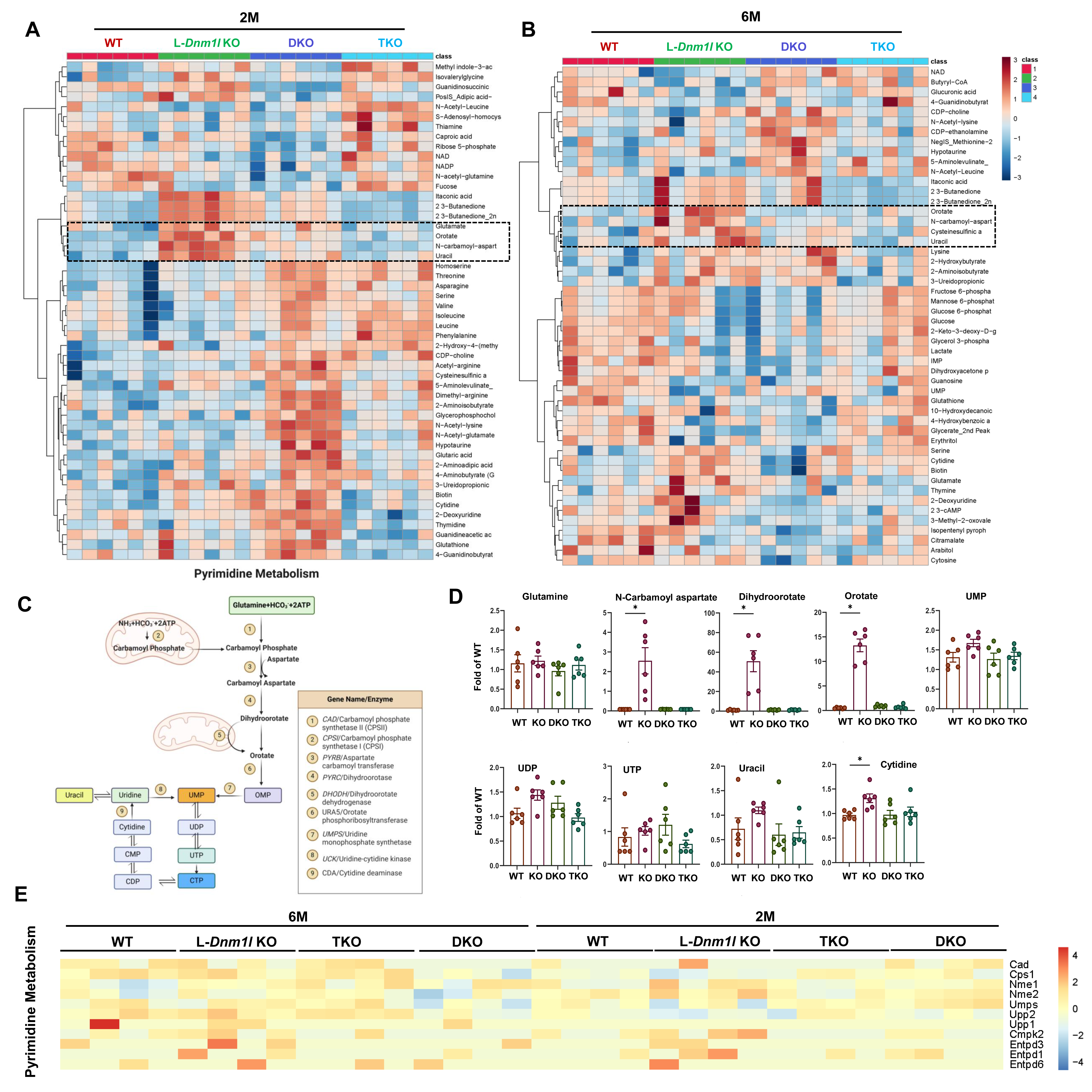
Increased pyrimidine synthesis and metabolism in the liver of L-*Dnm1l* KO but not TKO mice. Polar metabolites were extracted and subjected to LC-MS. Heatmap of targeted metabolomics analysis of indicated 2M (A) and 6M (B) mouse liver tissues (n=6). Boxed are pyrimidine related metabolites. (C) A summary graph of mitochondria and pyrimidine synthesis pathway. (D) Levels of key pyrimidine metabolites from untargeted metabolomics analysis (n=6). (E) Heatmap of pyrimidine metabolism genes from the RNA-seq analysis (n=6). All results are expressed as means± SE. *p<0.05, One-Way ANOVA analysis with Bonferroni’s post hoc test.

To better understand the broader mitochondrial dysfunction in L-*Dnm1l* KO mice, we next examined gene expression profiles associated with various mitochondrial components and functions. This analysis included genes related to complex I, complex II, complex III, complex IV, as well as mitochondrial cristae architecture, metabolic transporters, mitochondrial transcription, protein import, RNA/tRNA modification and processing, proteostasis and mitophagy, tRNA amino acid synthetases, ubiquinone (UQ) synthesis, mtRibosome subunits and assembly factors, mitochondrial translation, and mtDNA replication. In 2M L-*Dnm1l* KO mice, gene expression in these pathways generally showed a downward trend compared to WT mice but was largely corrected in TKO mice. However, the overall gene expression in above pathways was higher in 6M WT mice compared to 2M WT mice, and the differences between L-*Dnm1l* KO mice and WT mice were substantially reduced (**Supplemental Figures 4-8**), suggesting possible adaptive responses to mitochondrial dysfunction developed with age in L-*Dnm1l* KO mice.

### L-*Dnm1l* KO mice develop hepatocellular adenoma associated with increased tumor-related signaling pathways

To characterize the tumor type in L-*Dnm1l* KO mice, we performed histopathological analysis. H&E staining of tumor sections revealed the presence of hepatocellular adenoma in L-*Dnm1l* KO mice. The adenomas were poorly defined, with benign hepatocytes arranged in plates of regular thickness accompanied by infiltrated inflammatory cells, and lacked portal triads (**Figure 6A, white arrow**). Some tumor cells also showed marked steatosis, forming a steatotic adenoma indicating in deficiency of lipid metabolism (**Figure 6A, black arrow**). Reticulin is utilized to highlight the normal thickness of liver cell plates, excluding the diagnosis of well-differentiated HCC. Our findings indicate that certain liver tumors exhibited steatosis, intact reticulin fibers and lacked glypican 3 and nuclear β-catenin staining, which are characteristics of adenoma (**Fig. 6A, arrowhead**).

**Figure 6.**
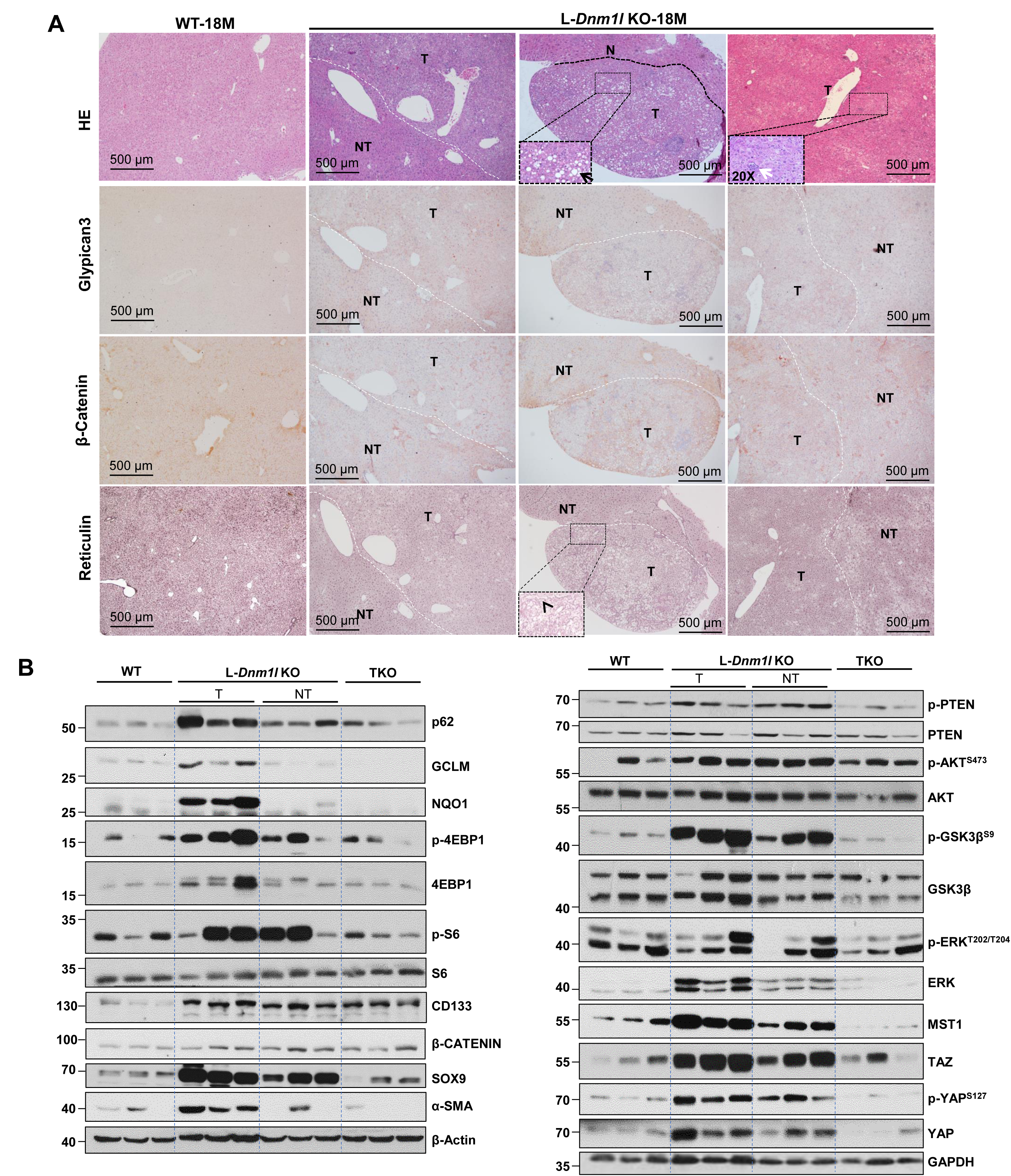
Hepatocellular adenoma in L-*Dnm1l* KO mice. (A) Representative H&E and immunohistochemistry staining of Glypican 3 and β-CATENIN as well as reticulin staining in 18M L-*Dnm1l* KO and matched WT mouse liver. White dotted line marks the boundary of tumor and non-tumor areas. Black arrow indicates hepatocellular adenoma with lipid accumulation, white arrow indicates infiltrated inflammatory cells, arrowhead indicates the reticulin positive staining. (B) Western blot analysis from total liver lysates or tumor and non-tumor tissues of indicated 18M mice. T: tumor, NT: non-tumor.

Increased levels of hepatic p62 have oncogenic effects by promoting the activation of NRF2 and mTORC1 pathways in the liver (Ni, Chao et al. 2019, Chao, Wang et al. 2022). The levels of hepatic p62, GCLM, and NQO1 (NRF2 target genes), along with p-S6 and p-4EBP1, were significantly higher in the tumors of L-*Dnm1l* KO mice compared to WT mice. However, these markers were reduced in L-*Dnm1l* KO non-tumor tissues and TKO mouse livers. Additionally, the levels of CD133 and SOX9 (markers associated with liver cancer stem cells), as well as various oncogenic factors such as β-CATENIN, p-PTEN, p-AKT, p-GSK3β, TAZ, and YAP, were markedly elevated in the tumors of L-*Dnm1l* KO mice compared to WT mice, L-*Dnm1l*

KO non-tumor tissues and TKO mouse livers. Interestingly, the major tumor suppressor mammalian STE20-like kinase 1 (MST1) and the downstream phosphorylated form of yes-associated protein 1 (p-YAP1) were increased in L-*Dnm1l* KO tumors compared to both WT and TKO mice. There were no significant changes in p-ERK levels, but an increase in α-SMA was observed in the L-*Dnm1l* KO tumors compared to WT and TKO mice (**Fig 6B**). Overall, these results indicate that hepatocellular adenoma, but not HCC, develops in L-*Dnm1l* KO mice. This development is associated with increase in certain tumor promoting and suppressing signaling pathways, all of which are absent in TKO mice.

### Increased DNA damage, senescence, and compensatory proliferation in the liver of L-*Dnm1l* KO but not TKO mice

DNA damage and cell proliferation play a crucial role in cancer development and progression. Results from the IHC staining showed an increased number of phosphorylated γ-H2AX and Ki67 positive cells in 2M L-*Dnm1l* KO mouse liver **(Fig. 7A-C)**. Intriguingly, the number of phosphorylated γ-H2AX and Ki67 positive hepatocytes was significantly decreased in 6M L-*Dnm1l* KO mice compared to 2M, suggesting a possible ongoing DNA lesion repair and adaptive process in these mice **(Fig. 7A-C)**. Consistent with this, p21 and p27, the target genes of tumor suppressor p53, whose induction leads to transcriptional downregulation of many cell cycle genes, as well as levels of PCNA and γ-H2AX, were upregulated in the tumors of 18M L-*Dnm1l* KO mice compared to WT mice **(Fig. 7D)**. There were higher number of γ-H2AX and Ki67 positive cells in the tumor areas than the adjacent normal hepatocytes (**Fig. 7E**). Furthermore, the number of β-Gal positive senescent cells increased in 18M L-*Dnm1l* KO and TKO mouse livers, although it only increased in the 2M TKO but not L-*Dnm1l* KO mouse livers **(Supplemental Fig. 9A-B).** Collectively, these data suggest that L-*Dnm1l* KO mice have increased DNA damage, and compensatory proliferation in the liver as early as 2M, which is absent in TKO mice.

**Figure 7.**
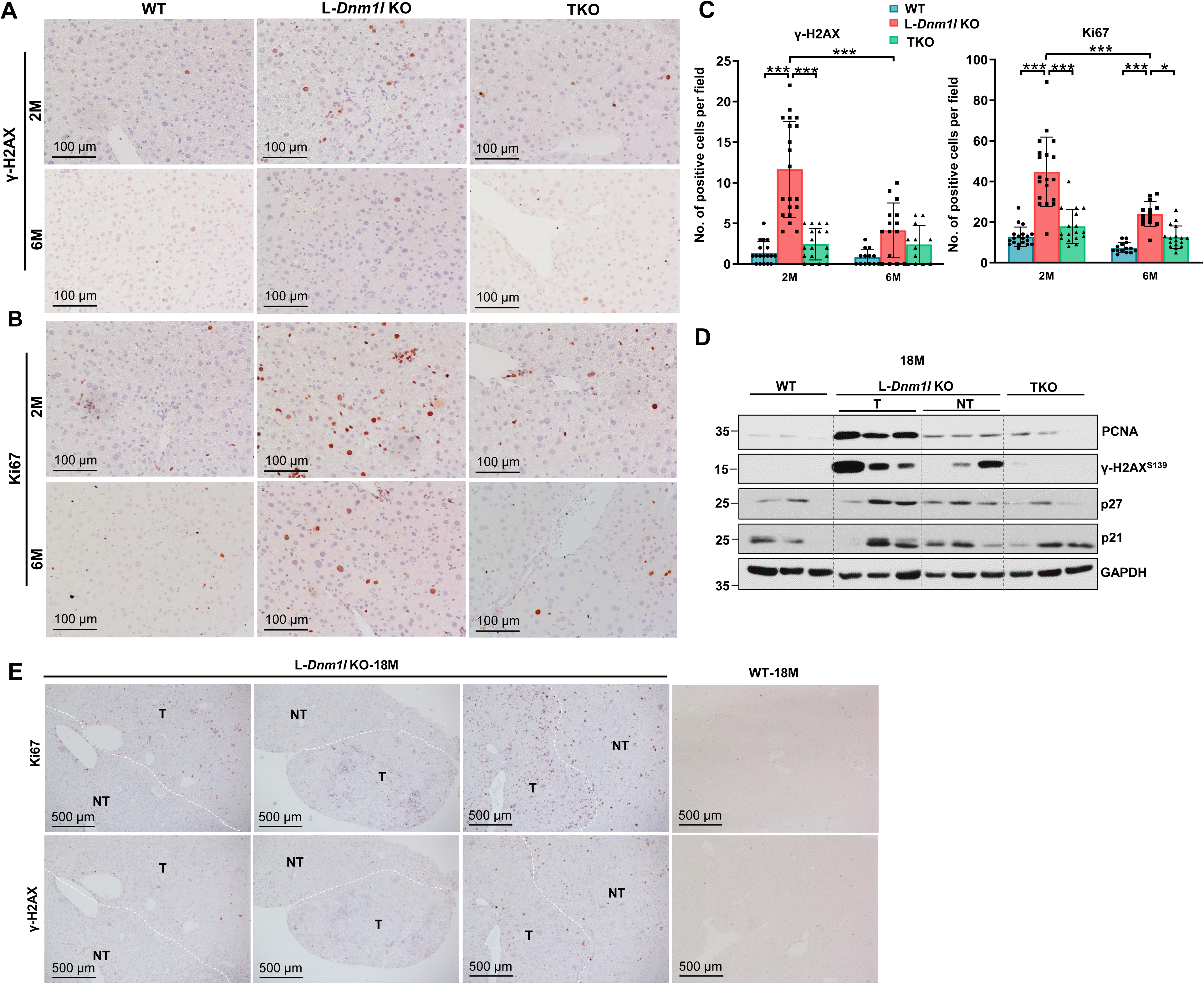
Increased DNA damage and proliferation in L-*Dnm1l* KO but not TKO mice. (A-B) Representative Immunohistochemistry staining of γ-H2AX and Ki67 of indicated mouse genotypes. (C) At least 5 fields were quantified from each mouse (n=3 mice). All results are expressed as means± SD. *p<0.05, **p<0.01, ***p<0.001; One-Way ANOVA analysis with Bonferroni’s post hoc test. (D) Western blot analysis from total liver lysates or tumor and non-tumor tissues of indicated 18M mice. (E) Representative Immunohistochemistry staining of γ-H2AX and Ki67 in 18M indicated genotyped mice. White dotted line marks the boundary of tumor and non-tumor areas. T: tumor, NT: non-tumor.

### Genetic restoration of mitochondrial stasis abolishes oncogene-driven liver cancer

To determine the role of mitochondria dynamics in oncogene-induced liver cancer, we used the sleeping beauty transposon and hydrodynamic tail vein injection (SB-HTVI) system to deliver c-*MYC* and the constitutively active *YAP* mutant *(YAP S127A)*, which has been characterized to induce HCC (Chen and Calvisi 2014). After 8 weeks following the hydrodynamic injection of *c-MYC/YAP*, visible tumor nodules appeared in mouse liver, with both tumor size and numbers, as well as serum ALT levels, increased in L-*Dnm1l* KO mice. Strikingly, hydrodynamic injection of *c-MYC/YAP* did not increase serum ALT levels in TKO mice, and only 2 out of 11 TKO mice developed visible tumors (**Fig. 8A-D**). H&E staining revealed typical HCC features of these tumors with increased nuclear cytoplasmic ratios. IHC staining showed increased Glypican 3 staining and Ki67 positive cells. These tumors were also positive for c-MYC and YAP proteins, confirming that they were driven by c-MYC/YAP (**Fig. 8E & Supplemental Fig. 10**). Notably, the levels of c-GAS, STING, TBK1 and IRF7, but not IRF3, were elevated in tumors of WT mice and further increased in L-*Dnm1l* KO mice. There are no significant changes of c-GAS, STING, TBK1 and IRF7 in non-tumor sections of TKO mice compared with WT mice (**Supplemental Fig. 10**).

**Figure 8.**
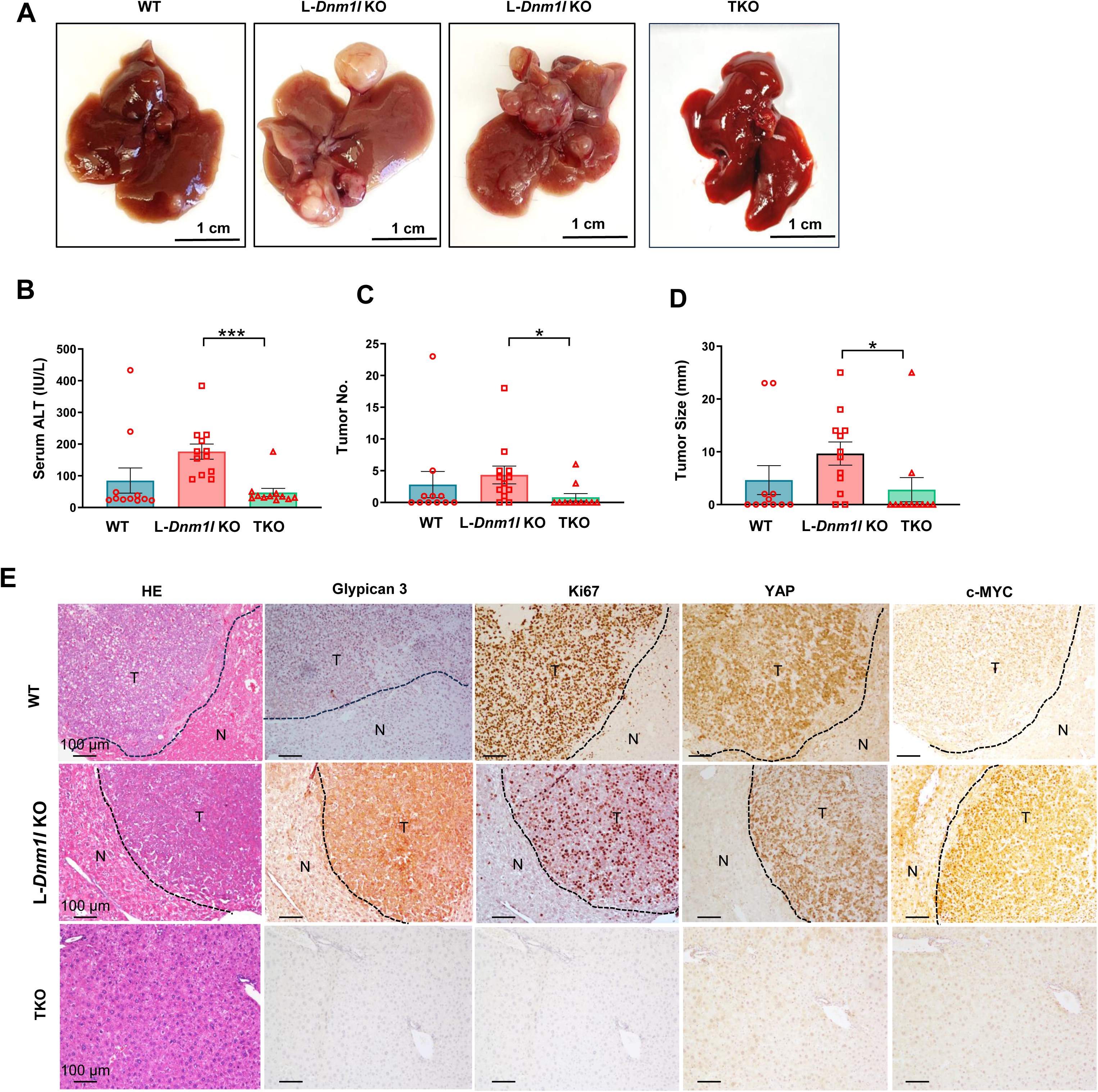
Increased oncogene-driven liver tumorigenesis in L-*Dnm1l* KO but not TKO mice. Sleeping beauty transposon (SB10) and c-MYC/YAP^S127A^ were delivered into 8-week-old male mice of indicated genotypes through hydrodynamic tail vein injection. Liver tissues and blood were collected 8 weeks post-injection. (A) Representative gross mouse liver images of indicated genotypes. (B) Serum ALT activities, (C) the number of tumors per mouse liver and (D) tumor size are quantified. All results are expressed as means± SEM (n=11-12). *p<0.05, Student’s t-test. (E) Representative H&E and IHC staining of indicated proteins in livers from mice with specific genotypes. Black dotted line marks the boundary of tumor and non-tumor areas. T: tumor, NT: non-tumor.

## Discussion

Mitochondrial morphological changes have been observed in various cancers and are considered a crucial metabolic factor that supports cancer cell growth and metastasis (Kashatus, Nascimento et al. 2015, Cheng, Kuo et al. 2016, Ma, Wang et al. 2020). Human HCC tissues are generally found to have higher levels of DRP1/MFN1 expression ratio, which was linked to a poor prognosis, and enhanced mitochondrial fission has been shown to promote HCC cell survival *in vitro* and *in vivo* via elevated ROS production (Huang, Zhan et al. 2016, Zhang, Li et al. 2020). These results led to a widely accepted notion that fragmented mitochondria promote the viability of cancer cells in non-permissive conditions. However, there is still no definite verdict on the actual mitochondrial size in human HCC tumor tissues. Indeed, direct evidence from two independent studies investigating mitochondrial length from paired HCC tumor-peritumor tissues through electron microscopy drew conflicting conclusions (Huang, Zhan et al. 2016, Li, Wang et al. 2020), indicating a strong heterogeneous metabolic feature of cancer cells in HCC. On the other hand, mitochondrial fusion was also found to support liver tumor growth (Li, Wang et al. 2020). Another study revealed that metabolic reprogramming via mitochondrial elongation is favorable for HCC cell survival and adaptation to energy stress (Li, Huang et al. 2017). Therefore, the role of mitochondrial architecture changes in liver cancer progression is complicated, and it seems fragmented and elongated mitochondria can be both pro-tumorigenic in the liver. In the present study, we found that deletion of either mitochondrial fission protein DRP1 or the two fusion proteins MFN1 and MFN2 in the liver resulted in spontaneous tumor formation in mice, and this phenotype can be rescued by restoration of mitochondrial dynamic balance through triple depletion of DRP1, MFN1 and MFN2. These findings suggest that mitochondria stasis is the key to preserve normal cellular functions in the liver and prevent tumorigenesis. Unlike previous reports showed that the removal of *Mfn1* or *Mfn2* promoted liver tumor formation either under aging-associated spontaneous conditions or triggered by carcinogen exposure (Hernandez-Alvarez, Sebastian et al. 2019, Li, Han et al. 2022, Wang, Tian et al. 2022), our data suggested that single KO of either *Mfn1* or *Mfn2* is not sufficient to cause age-related spontaneous tumor in the liver. It remains unclear whether differences in mouse strain background could account for such inconsistent observations, although it is highly likely that the loss of one MFN protein may be compensated by the other. In this study, we found a small portion (29% incidence) of L-*Mfn1/2* DKO mice developed tumors at 15-18M, however, the tumor burden and size in these mice were significantly lower than L-*Dnm1l* KO mice, suggesting that mitochondrial fission may play a more critical role than mitochondrial fusion in maintaining hepatocyte homeostasis and preventing tumorigenesis. As fragmented mitochondria are more readily removed by mitophagy, it is likely that L-*Dnm1l* KO mice have lower levels of mitophagy compared to L-*Mfn1/2* DKO mice, which would lead to increased accumulation of damaged mitochondria, elevated oxidative stress, DNA damage, genome instability and eventually, liver tumorigenesis.

One of the most intriguing findings of this study is the increased metabolism of pyrimidines observed in the livers of L-*Dnm1l* KO mice. Mitochondria are essential in pyrimidine metabolism, particularly in *de novo* pyrimidine synthesis, as the key enzyme DHODH catalyzing the conversion of dihydroorotate to orotate is located within the mitochondrial inner membrane (Bajzikova, Kovarova et al. 2019). It remains unclear whether increased megamitochondria formation affects the expression and activity of DHODH in pyrimidine biosynthesis. Nonetheless, an increase in the production of pyrimidine nucleotides, which serve as building blocks for DNA and RNA, may contribute to liver tumorigenesis in L-*Dnm1l* KO mice. Future studies are necessary to further investigate the role and mechanisms of megamitochondria in pyrimidine synthesis and metabolism during liver cancer development.

The activation of adaptive and innate immune responses is a key outcome of mitochondrial dysfunction, which has been implicated in tumor surveillance (Diefenbach and Raulet 2002). The cGAS-STING pathway is an evolutionarily conserved DNA-sensing defense mechanism that was initially identified for its role in protecting cells against viral infections (Decout, Katz et al. 2021). Accumulating evidence recently revealed a dual function of cGAS-STING signaling in both tumor suppression and promotion (Kwon and Bakhoum 2020). Acute activation of STING in early neoplastic cells leads to cell-cycle arrest via senescence-associated secretory phenotype (SASP). Experiment data indicated that cGAS or STING deficiency impairs senescence and SASP responses, accelerates spontaneous immortalization, and promotes tumor development (Dou, Ghosh et al. 2017, Gluck, Guey et al. 2017, Yang, Wang et al. 2017, Ranoa, Widau et al. 2019). However, chronic exposure to activated cGAS–STING signaling can lead cancer cells to lose downstream cell-cycle regulators, fostering tolerance. Additionally, sustained activation of inflammatory signaling in this state exhausts effector immune cells and recruits immunosuppressive cells, collectively creating an immune-suppressive tumor microenvironment that facilitates tumor survival and immune evasion (Toso, Revandkar et al. 2014, Bakhoum, Ngo et al. 2018, Georgilis, Klotz et al. 2018, Kwon and Bakhoum 2020). Here, we reported a remarkable upregulated cGAS-STING pathway in L-*Dnm1l* KO mouse liver, which is associated with increased number of senescent cells and elevated cell-cycle inhibitors, and these phenotypes become more evident in 15-18M tumor-bearing mice. Highly elevated cGAS/STING protein expression was observed in oncogene-induced tumor regions of L*-Dnm1l* KO mice. The role of cGAS-STING signaling as either an accelerator or suppressor of tumor progression after initiation is still unclear in our models. To clarity the causal role of this pathway in liver tumorigenesis, future experiments should focus on generating double KO mice for DRP1 and cGAS, as well as DRP1 and STING.

Another interesting finding from this study is that TKO mice, with reestablished mitochondrial fission/fusion stasis, were more resistant to oncogene-induced hepatic tumorigenesis than WT mice. This resistance is likely attributed to an enhanced SASP and reduced pyrimidine synthesis, as indicated by the presence of more senescent cells and lower pyrimidine metabolites in TKO mouse liver. The mitochondria in TKO mice are theoretically less active than those in WT mice due to the absence of both fission and fusion proteins, rendering them less responsive to external stress.

In conclusion, we found that deletion of mitochondrial fission protein DRP1 in the liver resulted in spontaneous tumor formation in mice. Mitochondrial dynamics and stasis are critical to maintaining hepatic mitochondrial homeostasis, pyrimidine metabolism and hepatocyte functions. L-*Dnm1l* KO mice had elevated pyrimidine metabolism and activated cGAS-STING signaling pathway, which is associated with increased DNA damage, senescence, and compensatory proliferation. These factors collectively accelerated tumorigenesis during aging or exposure to external oncogenes. This study lays the foundation for future drug development aimed at maintaining homeostasis of mitochondrial dynamics to preserve normal liver physiology and prevent cancer.

## Supporting information

Main text

