## Supplementary material for "Disruption of Mitochondrial Dynamics and Stasis Leads to Liver Injury and Tumorigenesis": Main text

**Supplemental Figure Legend**

**Supplemental Figure 1. Increased hepatic α-SMA levels in L-*Dnm1l* KO but not TKO mice.** Western blot analysis of total liver lysates of indicated mouse genotypes at 2M (A), 6M (B) and 12M (C).

**Supplemental Figure 2. L-*Dnm1l* KO mice but not DKO and TKO mice have increased activation of cGAS-STING-interferon pathway in the liver.**

(A) Principal component analysis (PCA) of RNA-seq dataset of 6M mice. (B) Heatmap of interferon and cGAS-STING pathway involved genes from the RNA-seq dataset. (C) Western blot analysis from total liver lysates or tumor and non-tumor tissues of indicated mice at 18M. (D) Western blot analysis of total liver lysates from mice of indicated genotypes at 18M. T: tumor, NT: non-tumor.

**Supplemental Figure 3. Liver cell type gene expression signature in WT, L-*Dnm1l* KO, DKO and TKO mice.**

(A) Pathway analysis of signature gene expression of hepatocytes, HSC and Kupffer cell/macrophage of RNA-seq dataset of 2M (A) and 6M (B) mice of indicated genotypes

**Supplemental Figure 4. Mitochondria complex assembly gene expression changes in L-*Dnm1l* KO, DKO and TKO mice.**

Heatmap of mitochondria complex assembly genes from the RNA-seq dataset of 2M and 6M mouse livers of indicated genotypes.

**Supplemental Figure 5. Mitochondria cristae architecture and metabolic transporter gene expression changes in L-*Dnm1l* KO, DKO and TKO mice.** Heatmap of mitochondria cristae architecture and metabolic transporter genes from the RNA-seq dataset of 2M and 6M mouse livers of indicated genotypes.

**Supplemental Figure 6. Mitochondria transcription and protein import gene expression changes in L-*Dnm1l* KO, DKO and TKO mice.**

Heatmap of mitochondria transcription and protein import genes from the RNA-seq dataset of 2M and 6M mouse livers of indicated genotypes.

**Supplemental Figure 7. Mitochondria mRNA/mRNA/rRNA modification and processing, proteostasis and mitophagy, TRNA amino acid synthetase and UQ synthesis gene expression changes in L-*Dnm1l* KO, DKO and TKO mice.** Heatmap of mitochondria mRNA/mRNA/rRNA modification and processing, proteostasis and mitophagy, TRNA amino acid synthetase and UQ synthesis genes from the RNA-seq dataset of 2M and 6M mouse livers of indicated genotypes.

**Supplemental Figure 8. Mitochondria Ribosome subunit assembly factors, mitochondria translation, and mtDNA replication gene expression changes in L-*Dnm1l* KO, DKO and TKO mice.**

Heatmap of mitochondria Ribosome subunit assembly factors, mitochondria translation, and mtDNA replication genes from the RNA-seq dataset of 2M and 6M mouse livers of indicated genotypes.

**Supplemental Figure 9. Liver cell senescence staining in L-*Dnm1l* KO, DKO and TKO mice.** (A-B) Representative images of β-Galactosidase (β-Gal) staining using liver cryosections of 2M and 18M mice of indicated genotypes (n=2 mice in each group). All results are expressed as means± SD. *p<0.05, **p<0.01, ***p<0.001; One-Way ANOVA analysis with Bonferroni's post hoc test.

**Supplemental Figure 10. Activated cGAS-STING pathway in oncogene-driven tumor sections of L-*Dnm1l* KO and WT mice.**

Sleeping beauty transposon (SB10) and *c-MYC/YAP*$^{S127A}$ were delivered into 8-week-old male mice of indicated genotypes through hydrodynamic tail vein injection. Liver tissues and blood were collected 8 weeks post-injection. Western blot analysis from tumor and non-tumor tissues of indicated mice. T: tumor, NT: non-tumor.

#### Supplemental Figure 1

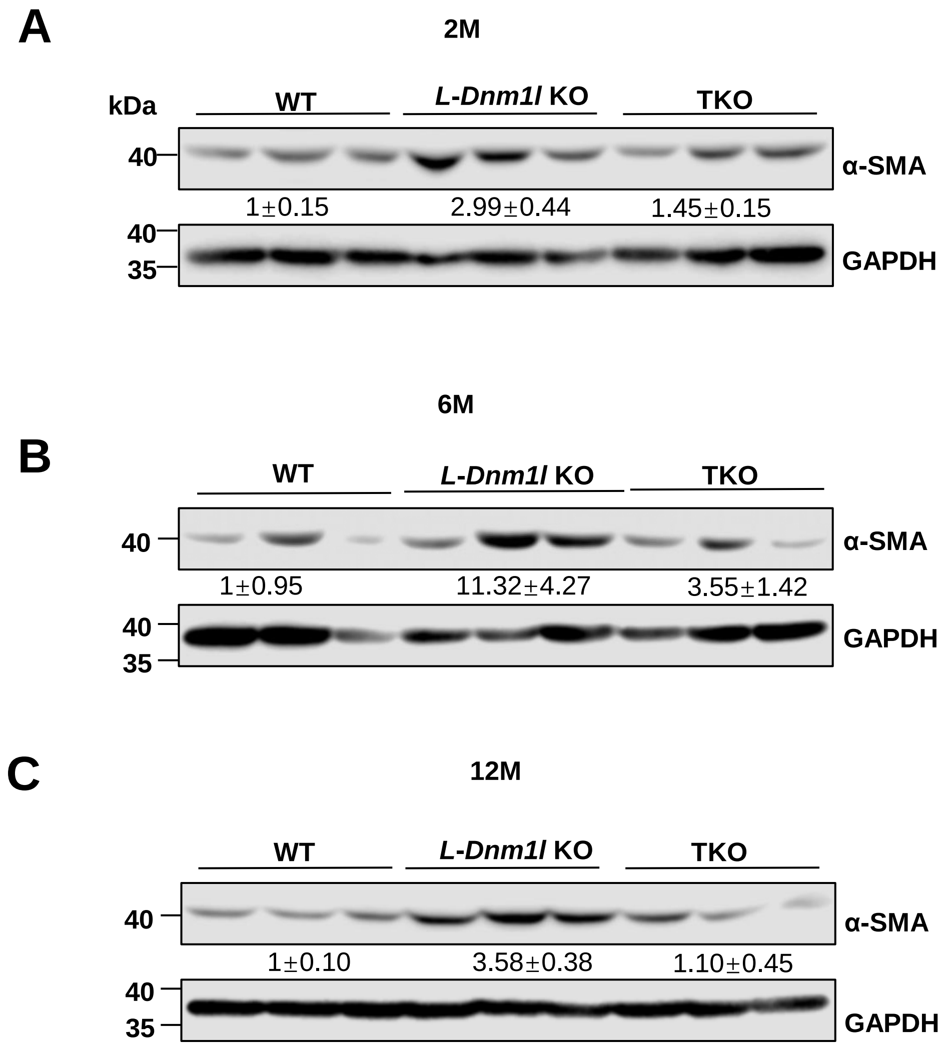

#### Supplemental Figure 2

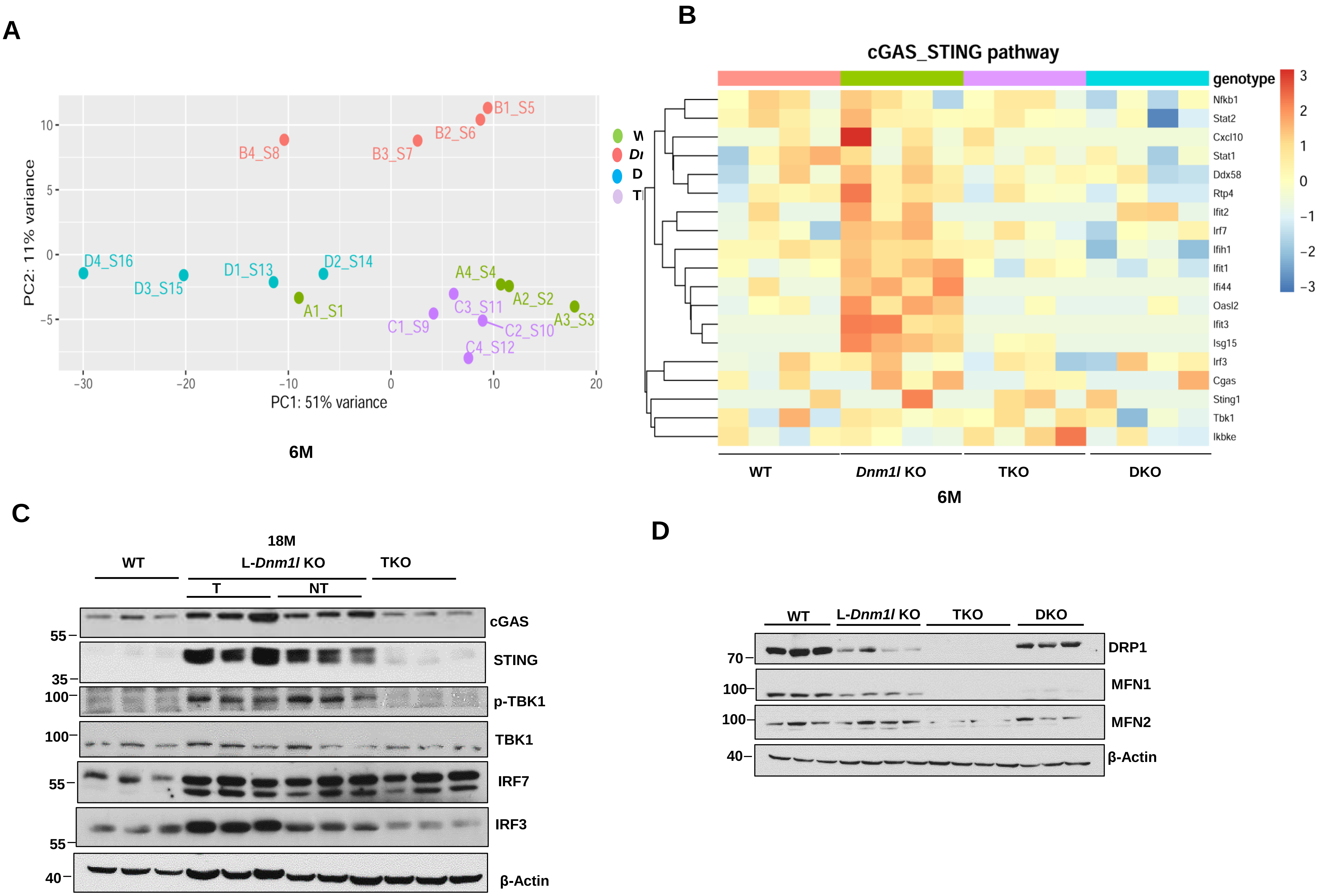

Supplemental Figure 3

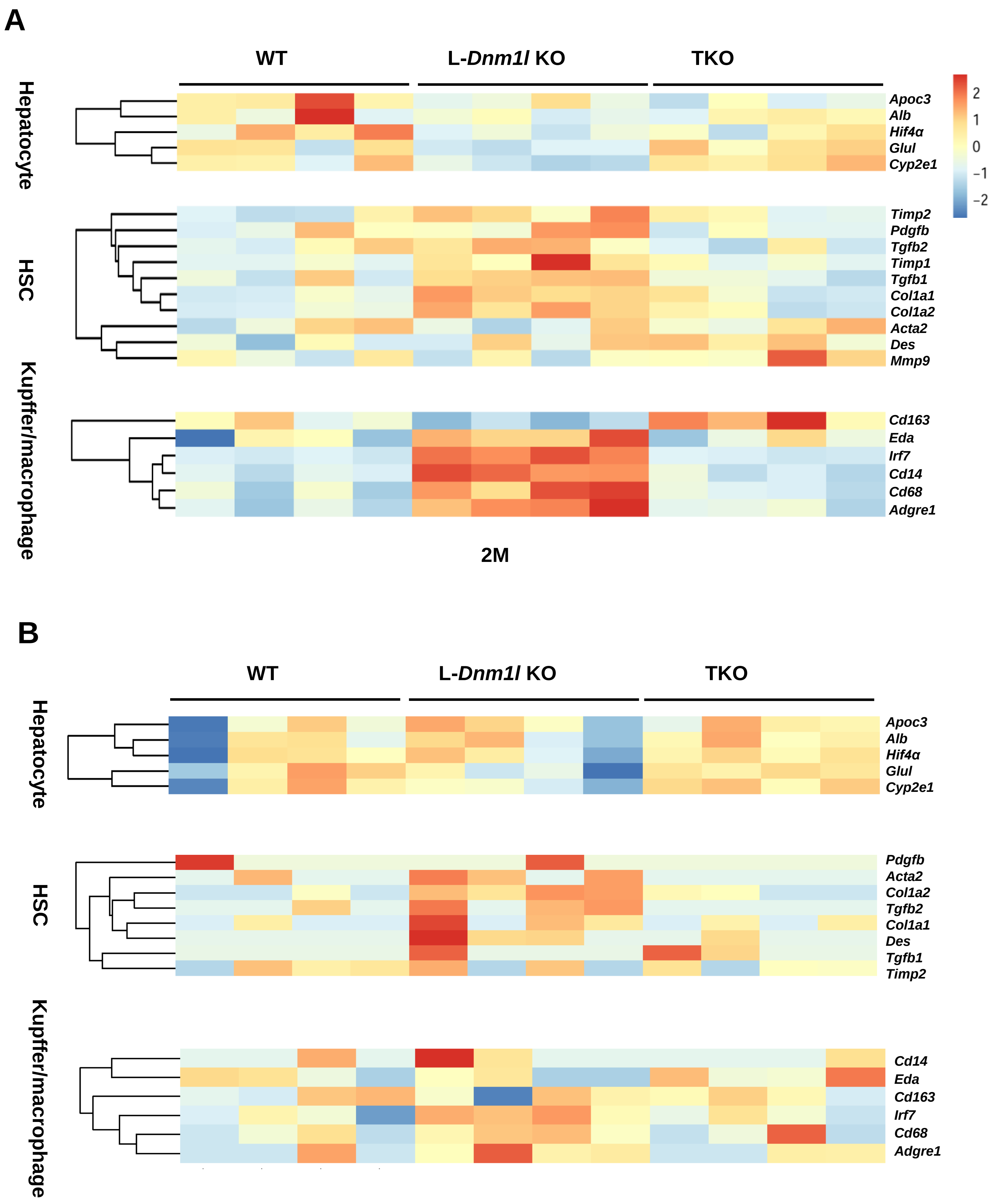

**Supplemental Figure 4**

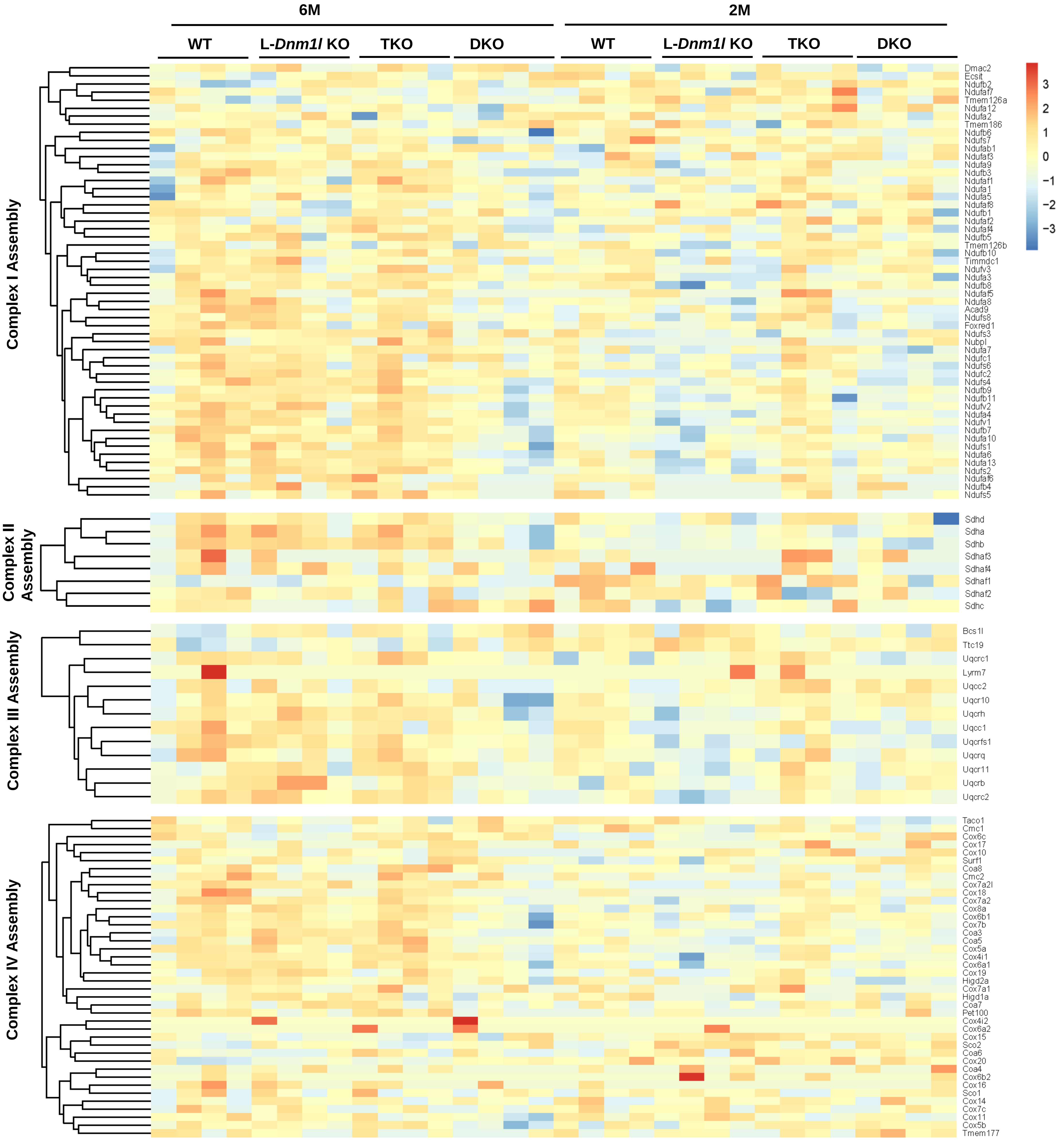

#### Supplemental Figure 5

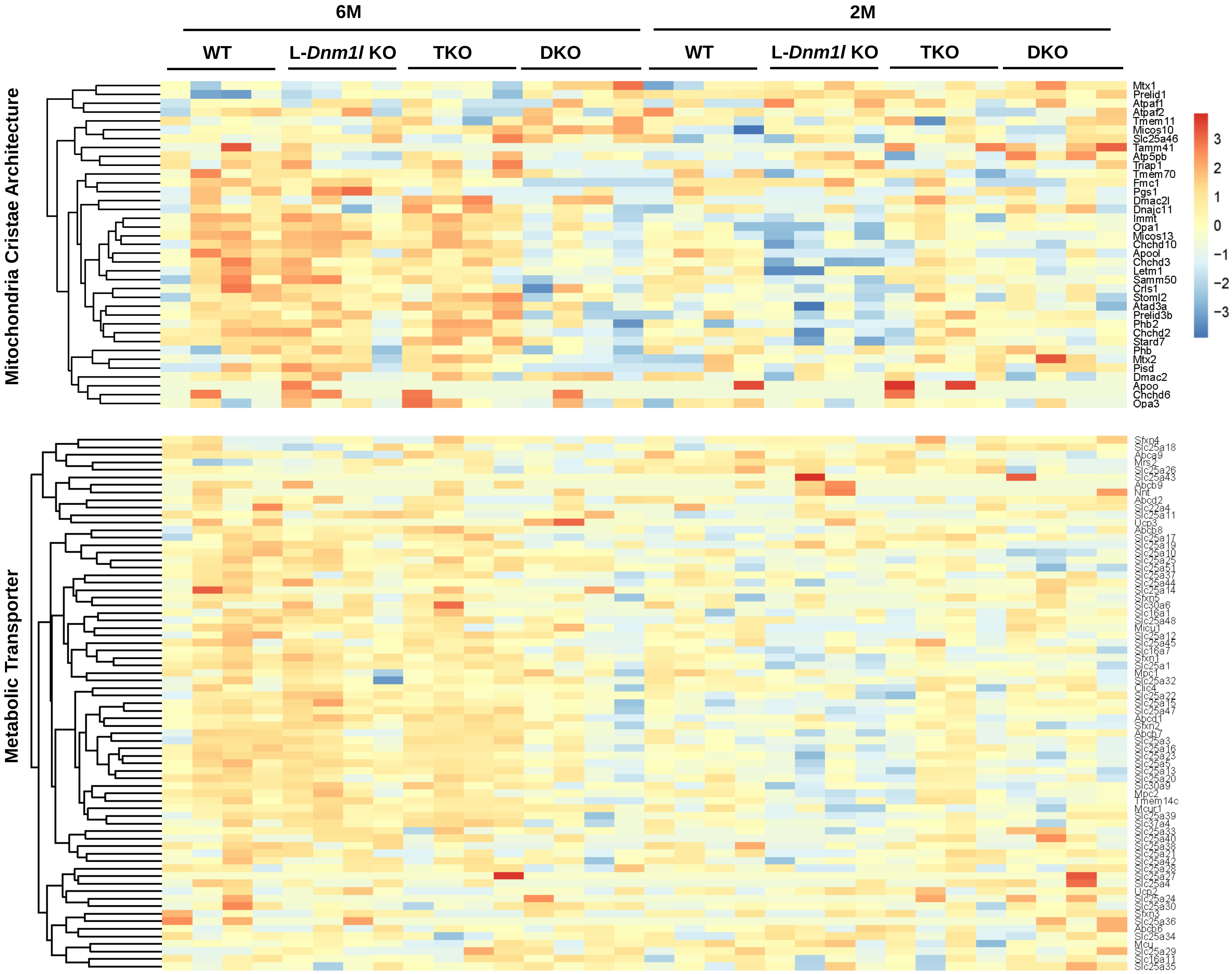

#### Supplemental Figure 6

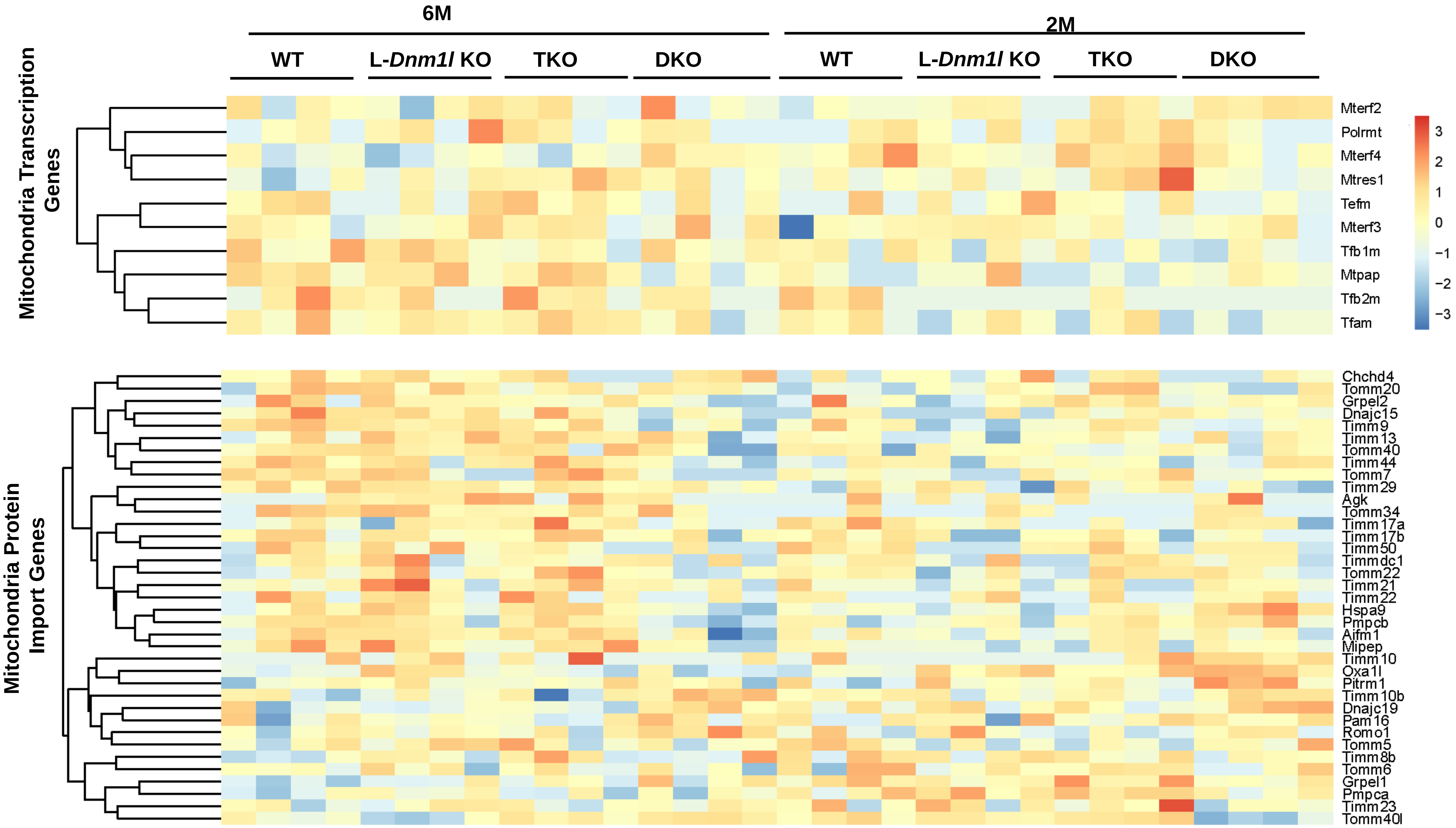

### Supplemental Figure 7

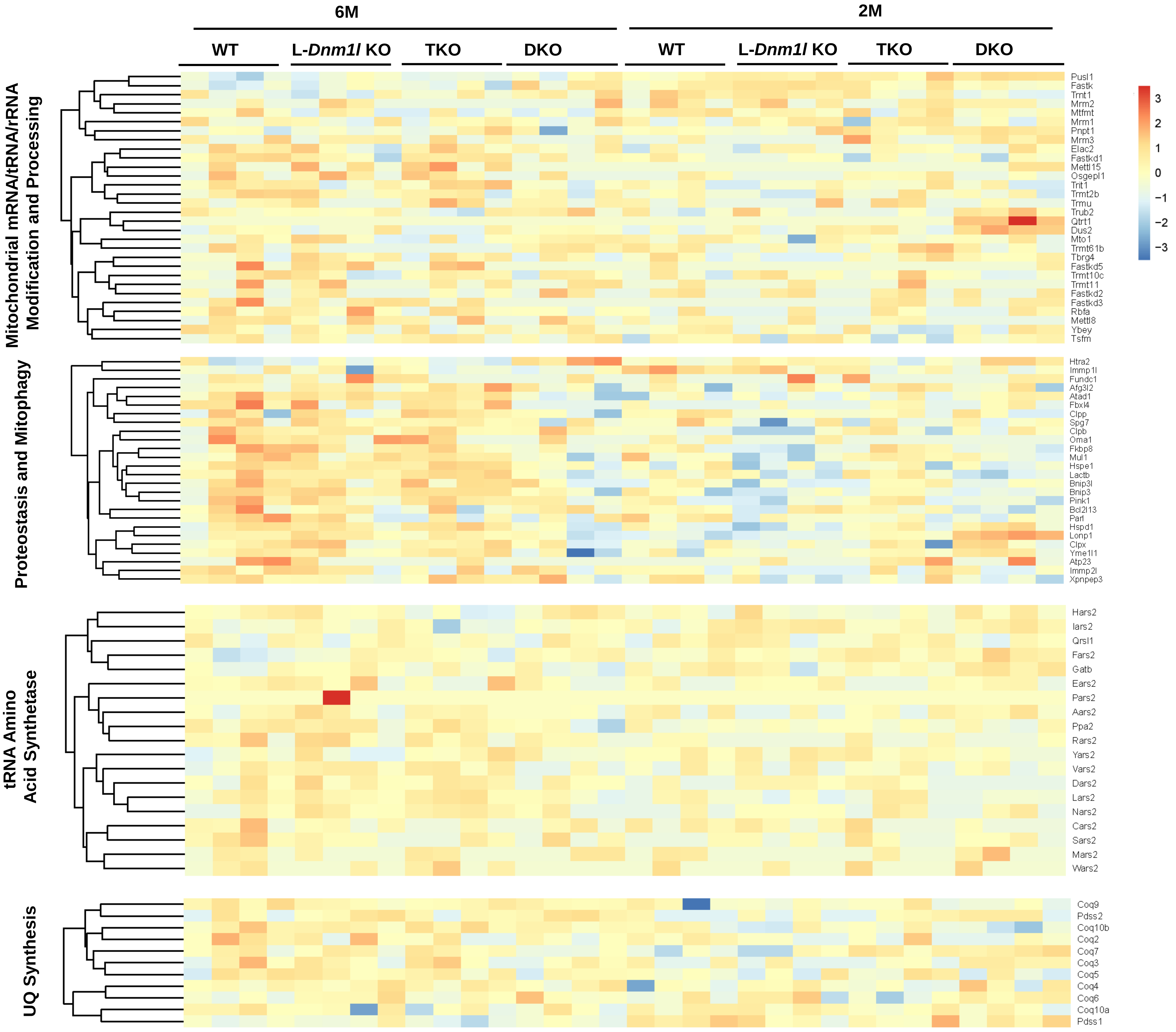

### Supplemental Figure 8

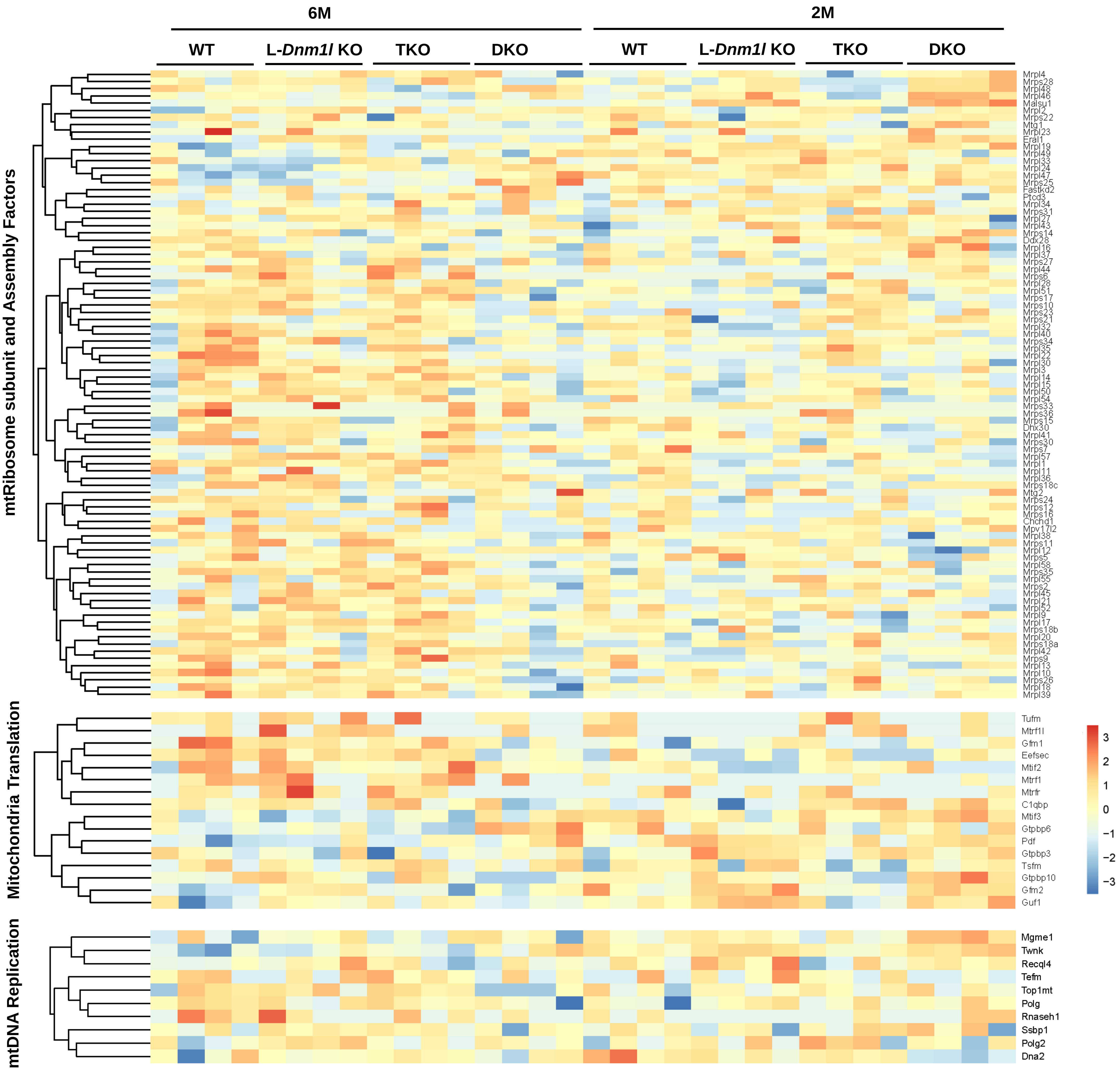

### Supplemental Figure 9

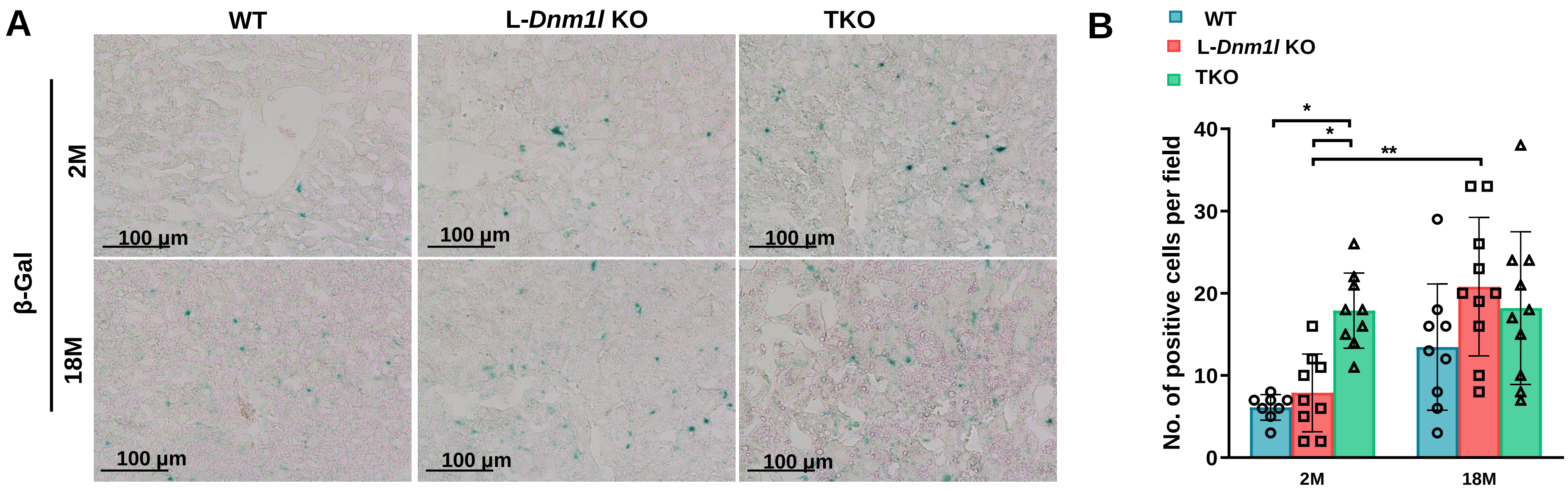

### Supplemental Figure 10

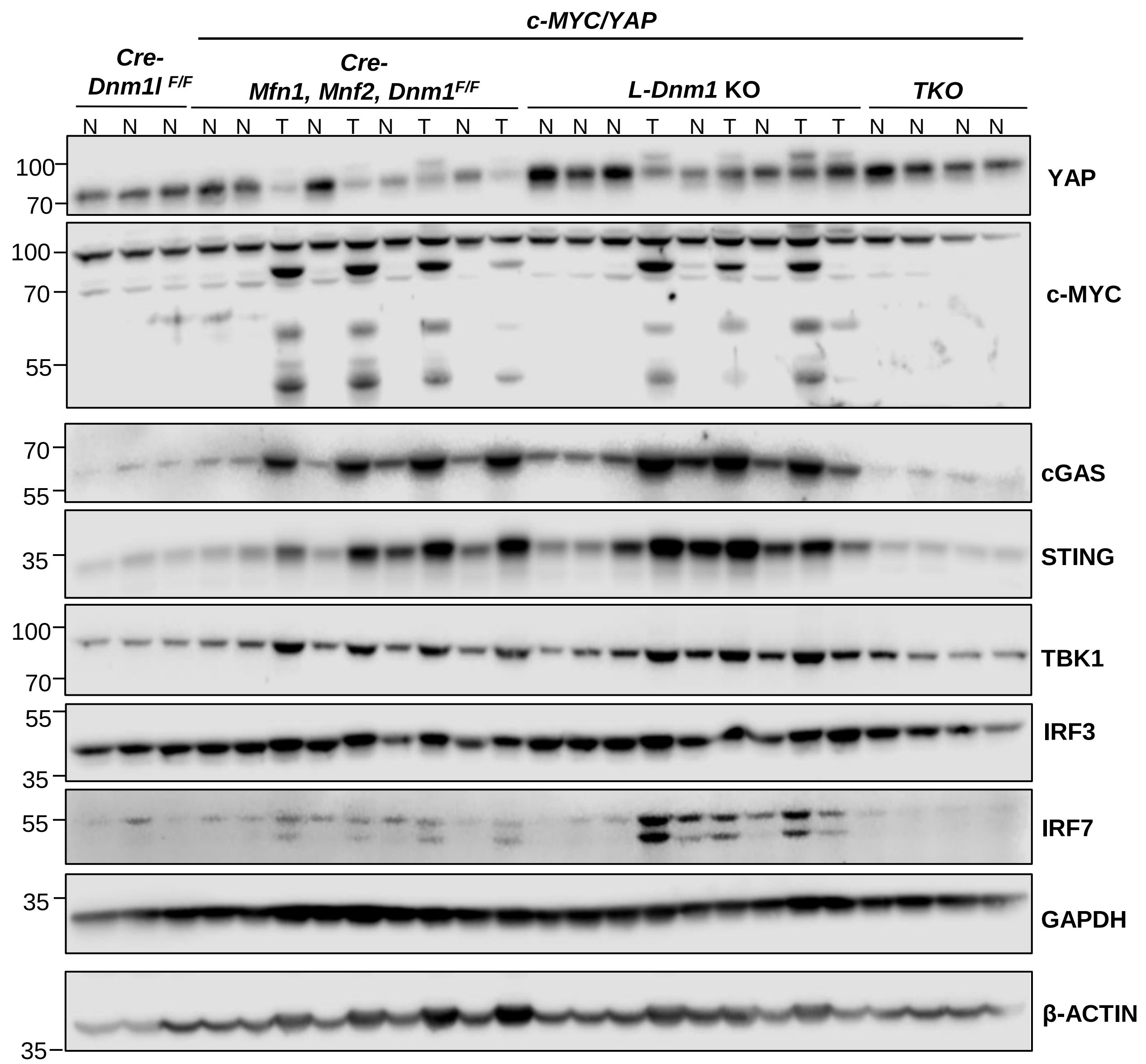
